# Non-canonical reductive nitrous oxide production pathways in a seasonally stratified lake basin

**DOI:** 10.1101/2025.08.15.670496

**Authors:** Teresa Einzmann, Moritz F. Lehmann, Jakob Zopfi, Claudia Frey

## Abstract

Nitrous oxide (N_2_O), a potent greenhouse gas and ozone-depleting agent, was detected at high concentrations in the anoxic bottom-waters of a monomictic eutrophic lake basin in Switzerland (Lake Lugano). The observed high site-specific nitrogen (N) isotope preference (SP), was inconsistent with bacterial denitrification, which typically exhibits low SP, thereby challenging its role as the primary N_2_O source. This pointed to chemo-denitrification and/or fungal denitrification, both characterized by high SP, as possible alternative pathways. We conducted incubation experiments with sediment and bottom-water samples to assess N_2_O production and reduction dynamics and associated natural-abundance stable isotope signatures. We demonstrate that N_2_O accumulation predominantly originated from sedimentary production, and that elevated SP values in the bottom water reflected fractional bacterial N_2_O reduction. Using an isotope mass balance mixing model, we identified bacterial denitrification as the dominant sedimentary process (∼75%), followed by chemo-denitrification (∼20%), and fungal denitrification (∼5%). Additional ^15^N tracer incubation experiments, combined with selective inhibitors to quantify isolated pathways, confirmed model-estimated contributions. These findings validate the use of literature-based SP values in mixing models, and provide evidence for non-canonical N_2_O production via chemo- and fungal denitrification, highlighting the need to broaden our understanding of N_2_O cycling in lakes beyond classical bacterial pathways.

**SYNOPSIS:** Eutrophic lake conditions significantly enhance nitrous oxide production through diverse microbial processes, including non-traditional pathways beyond classical nitrification and denitrification. This study highlights the importance of incorporating chemo- and fungal denitrification when investigating aquatic nitrous oxide cycling.

**GRAPHICAL ABSTRACT:** 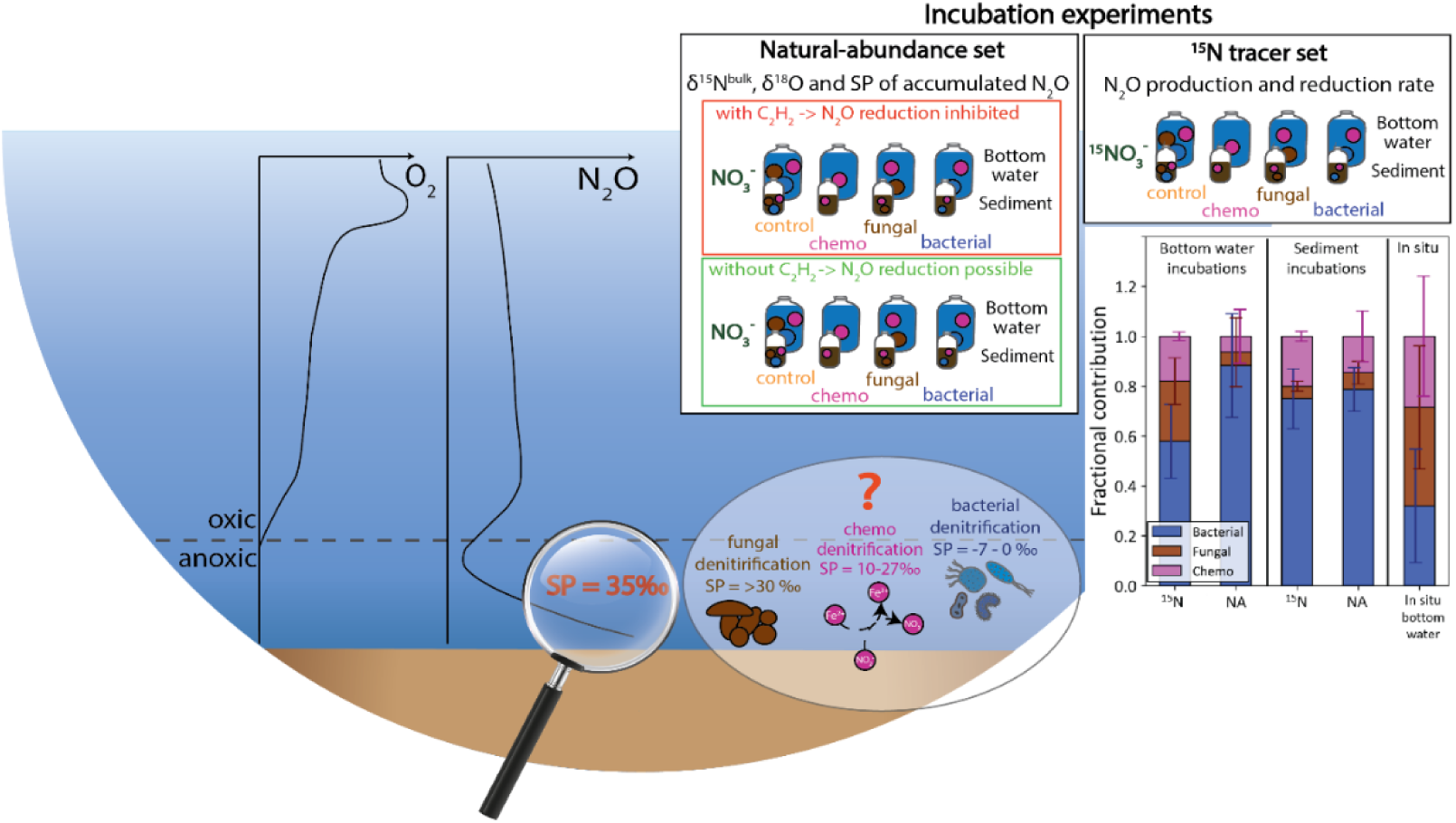

## 1 INTRODUCTION

Nitrous oxide (N_2_O) is a potent greenhouse gas with a warming potential up to 300 times greater than that of carbon dioxide (CO_2_) over a 100-year timescale, and it is the primary agent responsible for ozone depletion.^1^ Since the pre-industrial era, atmospheric N_2_O concentrations have increased from 270 ppb in 1750 to 336 ppb in 2022, with a record growth rate of 1.38 ppb per year in 2021.^2^ A major driver of this trend is the intensification of agricultural fertilization leading to increased nitrogen (N) loading into ecosystems, and results in elevated N_2_O emissions from terrestrial and aquatic environments.^3,4^ Despite evidence of rising N_2_O emissions from inland waters,^5^ some freshwater environments act as sinks for atmospheric N_2_O.^6,7^ Understanding the dual role of lacustrine systems in both N_2_O production and removal is crucial for predicting their response to anthropogenic nutrient inputs.

In aquatic systems, N_2_O is primarily produced through three key microbial pathways: nitrification, nitrifier-denitrification and denitrification. Nitrification, an aerobic process, involves the oxidation of ammonia (NH₃) or ammonium (NH_4_^+^), the latter being the dominant species of NH_4_^+^/NH_3_ in the pH range of most natural aquatic system, to nitrate (NO_3_^-^) via nitrite (NO₂⁻), with N_2_O released as a by-product. Under low-oxygen conditions, nitrifiers may simultaneously reduce NO_2_^-^ to N_2_O and dinitrogen (N_2_), a process known as nitrifier-denitrification. In contrast, denitrification is an anaerobic process where bacteria reduce NO_3_^-^ to N_2_, with N_2_O formed as an intermediate. Dissimilatory nitrate reduction to ammonium (DNRA) is another anaerobic pathway, which converts NO_3_^-^ to NH_4_ ^+^ via NO_2_^-^, and can also lead to N_2_O production. However the underlying enzymatic mechanisms are not fully understood.^8,9^

Of these processes, denitrification is the only known biological sink for N_2_O, especially under oxygen-depleted conditions. However, many denitrifying bacteria lack the complete gene set necessary to reduce N_2_O to N_2_, resulting in a truncated denitrification pathway.^10,11^ Moreover, the enzymatic reactions involved in denitrification vary in their sensitivities to oxygen, with nitrous oxide reductases (NOS) being particularly oxygen-sensitive.^12^ As a result, N_2_O often accumulates under suboxic conditions while complete denitrification dominates in the anoxic part of the water column. Thus, oceanic oxygen-deficient zones (ODZ), for example, enhance N_2_O consumption, whereas conditions along suboxic-anoxic interfaces favour N_2_O production, leading to higher N_2_O yields due to incomplete denitrification.^13–15^

Recently, alternative N_2_O production pathways have been identified, including denitrification by fungi and chemo-denitrification, an abiotic process involving reactions between ferrous iron (Fe(II)) and nitrite or nitrate in coastal sediments and ferruginous waters.^16–18^ Denitrifying fungi, which lack the gene encoding NOS, produce N_2_O as a terminal product, making their role in N_2_O production ecologically relevant.^19^ Fungal denitrification is a recognised source of N_2_O in terrestrial and coastal environments,^16,17,20,21^ and has also been implicated in the open ocean, where 18-22% of N_2_O at the oxic-anoxic interface of an ODZ may be of fungal origin.^22^ Despite growing evidence for both fungal and abiotic N_2_O production, the role of these alternative N_2_O production pathways in freshwater lakes remains underexplored. This, in turn, suggests that microbial denitrification may have been overestimated as the dominant source of N_2_O in lacustrine environments.

Stable N and O isotopic analysis of N_2_O is widely used to differentiate among N_2_O production pathways. Isotopic signatures depend the kinetic isotope effects (ε), which reflect the preferential use of the lighter isotopes (e.g., ^14^N versus ^15^N) during enzymatic or abiotic reactions, and on the isotopic composition of precursor substrates. Moreover, the intramolecular distribution of ^15^N between the central (α) and outer (β) N atoms, expressed as site preference (SP = δ¹⁵Nα – δ¹⁵Nβ), provides insight into the formation mechanisms.^23,24^ Characteristic SP ranges have been reported for different N_2_O production pathways: ∼-8 to 4‰ for incomplete bacterial denitrification, ∼30 to 39‰ for fungal denitrification, and ∼10 to 27‰ for chemo-denitrification.^25–27^ However, the use of SP for differentiating N_2_O production pathways is not always straightforward. Hybrid N_2_O formation, where N atoms originate from different substrates, can yield variable SP values (Kelly et al. 2024). Additionally, flavohemoglobin (fhp), a nitric-oxide (NO) reducing, detoxifying enzyme common in bacterial denitrifiers, has been shown to produce N_2_O with elevated SP values (∼11‰) compared to respiratory nitric oxide reductases (NOR).^28^ Furthermore, N_2_O reduction increases SP,^29^ potentially obscuring the SP signature of the production process. To address this, combining SP with δ^18^O-N_2_O measurements has proven effective in disentangling N_2_O production and reduction processes.^16,30^

In this study, we investigate reductive N_2_O production pathways in the bottom water and surface sediment in the South Basin (SB) of Lake Lugano (Switzerland), where N_2_O concentrations as high as 900 nM have been observed during summer stratification.^31^ High sediment-to-water N_2_O fluxes have been observed year-round,^32^ but previous studies have attributed N_2_O production in the sediment solely to bacterial denitrification, potentially overlooking alternative pathways.

We performed incubation experiments with bottom water and surface sediments to determine the relative contributions of bacterial, fungal and chemo-denitrification to N_2_O production/ accumulation in deep-hypolimnetic waters of Lake Lugano during summer stratification. Incubation experiments with ^15^N-NO_3_^-^, combined with pathway-specific inhibitors, enabled quantification of N_2_O production and consumption rates. Parallel natural-abundance (NA) incubations, conducted under the same inhibitory conditions, were used for isotopic signature identification of N_2_O. Dual-isotope mapping and a Bayesian isotope mixing model were applied to the NA-incubation data to apportion contributions from individual N_2_O formation pathways. The ^15^N-incubations further served to validate the reliability of using literature-derived isotope endmembers for N_2_O source attribution in freshwater systems.

## 2 MATERIALS AND METHODS

### 2.1 Study site and sampling

The study was conducted in the monomictic, eutrophic South Basin (SB) of Lake Lugano, located at the Swiss/Italian border. The basin stratifies seasonally from June to January and has a maximum depth of 95 m.^32,33^ Sampling was carried out in June and August 2023 at the basin’s deepest point (8°53’37”E, 45°57’31”N). A conductivity-temperature-depth profiler (CTD) (SBE 19plus V2 SeaCAT Profiler CTD, Seabird) was used to measure temperature, oxygen concentrations and turbidity in the water column. The first sampling campaign in June focussed on assessing thermal stratification and bottom-water N_2_O accumulation, while the August sampling targeted the collection of bottom-water and surface sediments for incubation experiments.

Duplicate samples for N_2_O concentration and isotopic analysis were collected at 5-10 m depth intervals using 5 L Niskin bottles. After overflow, samples were transferred into acid-washed, muffled (6 h at 450 °C) 160 mL glass serum bottles via gas-tight Tygon tubing. A 10 mL headspace was created, and 3 mL of 10 M NaOH was added as a preservative. Bottles were sealed with a grey butyl rubber septum (hollow stoppers grey, 20 mm, Chromatographie Service GmbH), crimped, and stored upside down at room temperature in the dark until isotope-ratio mass spectrometry (IRMS) analysis.

For NO_3_^-^ and NO_2_^-^ analysis, samples were filtered (0.2 µm pore-size, Filtropur S, Sarstedt) and frozen at -20 °C. Samples for Fe(II) analysis were not filtered and were fixed with sulfamic acid (40 mM final concentration).

For incubation experiments with unfiltered bottom water, water was collected at 93 m depth directly into muffled 2 L Schott bottles and sealed with a rubber stopper to avoid oxygen contamination. For filtered bottom-water incubations, water was passed through a Sterivex filter (Millipore, membrane material, 0.2 µm) using a peristaltic pump. For sediment slurry incubations, three sediment cores (diameter: 6 cm) were collected using a custom-made gravity corer, transported back to the laboratory and stored in the cold room until further processing within 24 hours. For slurry incubations, the top 5 cm of each core were combined and homogenised and bottom water was filtered as described above into acid-washed (10% HCl) and DI water-rinsed (5x times) plastic canisters. All samples were transported to the laboratory at 4 °C in the dark.

### 2.2 Chemical and stable isotope analyses

Concentrations of NO_3_^-^ were quantified by ion chromatography (940 Professional IC Vario, Methrom), while NO_2_^-^ was determined using standard spectrophotometric methods involving sulfanilamide and the Griess reagent (naphtal-ethylenediamine dihydrochloride).^34^ Concentrations of Fe^2+^ in sulfamic-acid-preserved samples were determined spectrophotometrically using the ferrozine assay.^35^

For N_2_O concentration and isotopic analysis, serum bottles were purged with helium (He) as carrier gas for 38 min. The N_2_O was trapped in liquid N_2_ following H_2_O removal using an ethanol trap (-60 °C) and a Mg(ClO_4_)_2_ trap; CO_2_ was removed with Ascarite. Remaining traces of CO_2_ and sample N_2_O were separated using a gas chromatograph (GC). Masses 44, 45, 46 (and their respective isotope ratios 45/44, 46/44) of the purified N_2_O, as well as the NO^+^ fragment ions at masses 30 and 31 (and isotope ratio 31/30), were analyzed by GC-IRMS (Delta V Plus, Thermo), with reference gas injection and flow control handled via a Conflo IV interface. The N_2_O isotope ratios were calibrated against reference N_2_O gas (≥ 99.9986%, Messer), aligned to the Tokyo Institute of Technology scale (Mohn et al. 2012) for both bulk and site- specific isotopic composition. Ratios of m/z 45/44, 46/44, and 31/30 signals were converted to δ^15^N-N_2_O (referenced to AIR), δ^18^O-N_2_O (referenced to Vienna Standard Mean Ocean Water, V-SMOW), and site-specific δ^15^N_α_ and δ^15^N_β_-N_2_O using three calibration standards (mixtures of N_2_O in synthetic air; CA06261, 53504, CA08214) and two scrambling factors (gamma and kappa) according to Frame and Casciotti (2010). N_2_O isotopocule corrections were performed using the *pyisotopomer* Python package (Kelly et al. 2023).

N_2_O concentrations were calculated by calibrating the total peak areas against standards of known concentrations, prepared by concerting 20 nmol NO_3_^-^ standards to N_2_O using the denitrifier method (see below).

The isotope ratios of NO_3_^-^ (N and O) were measured using the denitrifier method,^36,37^ in which NO_3_^-^ was converted to N_2_O by *Pseudomonas chlororaphis subsp. aureofaciens* (ATCC 13985) and subsequently analysed by GC-IRMS as above. Nitrate isotope measurements were calibrated using the international NO_3_^-^ standards IAEA-N3 (δ^15^N = 4.7‰, δ^18^O = 25.6) and USGS34 (δ^15^N = -1.8‰, δ^18^O = -27.9‰), and an in-house standard (UBN-1; δ^15^N = 14.15‰, δ^18^O = 25.7‰).

Isotope ratios of N_2_ were determined with a GasBench II coupled to an IRMS (Delta V Plus, Thermo). Air N_2_ was used as both calibration and drift-correction standard for δ^15^N-N_2_.

### 2.3 Incubation experiments

Two sets of experiments were conducted: one with ^15^N-labelled nitrate to measure N_2_O production and consumption rates, and another with NA nitrate additions to examine shifts in δ^18^O and SP of the produced N_2_O in association with specific processes. In both experimental sets, four treatments with specific inhibitors were applied to selectively suppress different N_2_O production pathways (Table S1 and S2).

The four treatments were as follows: 1. *Control* - with no inhibitors added; 2. *Bacterial* - fungal inhibitor added, allowing N_2_O accumulation only from incomplete bacterial or chemo- denitrification; 3. *Fungal* - bacterial inhibitor added to allow N_2_O production only via fungal or chemo-denitrification; 4. *Chemo* - both fungal and bacterial inhibitors added to supress biological N_2_O production completely, leaving only chemo-denitrification. Although chemo-denitrification may occur in all treatments to some degree, treatment 2 and 3 are referred to as *bacterial* and *fungal* for clarity. The occurrence of DNRA and its potential contribution to N_2_O production was not considered in our incubation experiments; however the importance of DNRA in the SB of Lake Lugano was investigated in previous studies, reporting varying contributions below 12%^38^ and up to 50%^39^ to NO_3_^-^ reduction compared to bacterial denitrification.

Streptomycin (1.29 mM final concentration) and cycloheximide (5.33 mM) were used as bacterial and fungal inhibitors, respectively, following concentrations previously shown to achieve up to 90% inhibition.^16,40^ A 24 h pre-incubation period allowed the inhibitors to take effect. Afterward, all vials were He-purged for 30 minutes to remove O_2_ and N_2_O.

To ensure sufficient Fe(II) availability for chemo-denitrification (in case in situ Fe(II) had already been oxidised before the start of the incubations), ferrous iron was added as anoxic FeSO_4_ solution to a final concentration of 2 µM, reflecting in situ lake conditions. No Fe(II) was added to filtered bottom-water samples, which served as abiotic controls. In sediment incubations, Fe(II) was assumed to be naturally available based on previously determined porewater and solid-phase profiles.^41^ Oxygen concentrations were monitored in separate incubation bottles per treatment using non-invasive optical trace-range O_2_ sensor spots (Pyrocience, Firesting, detection limit 0.005% O_2_).

### 15N-incubation experiments

^15^N-incubations were conducted to determine potential N_2_O production rates following the addition of ^15^N-labelled KNO_3_ (≥98 atom % ^15^N, Cambridge Isotopes). For bottom-water incubations (^15N^BW), 18 ml of inhibitor-treated bottom water was filled into muffled 20 mL glass serum vials, leaving a 2 mL headspace. For sediment incubations (^15N^Sed), 2 g of mixed sediment (top 5 cm) was transferred into 20 ml vials, and 8 ml of treated, filtered bottom water was added. All vials were crimp-sealed and shaken vigorously. After 24 h-pre-incubation and subsequent He-purging (30 min.), natural-abundance N_2_O was injected into each vial to achieve a background concentration of ∼50 nM. This ensured sufficient total N_2_O for reliable isotopic measurements by IRMS and allowed detection of ^15^N-labelled N_2_O as it accumulated during the incubation. Experiments comprised a total of eight treatments: 1) ^15N^BW-Control, 2) ^15N^BW-Bacterial, 3) ^15N^BW-Fungal, 4) ^15N^BW-Chemo, 5) ^15N^Sed-Control, 6) ^15N^Sed- Bacterial, 7) ^15N^Sed-Fungal, 8) ^15N^Sed-Chemo (Table S1). A filtered bottom-water treatment without Fe(II) served as negative abiotic control.

Each vial received ^15^N-NO_3_^-^ to a final concentration of 10 µM, and was incubated in the dark at 7 °C. For each treatment, 15 replicate vials were prepared. At designated time points (0, 6, 12, 16, 24 h for ^15N^BW; 0, 1, 2, 4, 6 h for ^15N^Sed) three vials per time point were sacrificed by adding 200 µL of 10 M NaOH.

For N_2_O reduction rate determination, 6 ml of liquid was subsampled from each vial prior to NaOH addition, transferred to He-purged exetainers, and subsequently analysed for δ^15^N_2_.

Two assumptions were made for calculating N_2_O production: 1) chemo-denitrification occurs across all treatments and must be subtracted from total N_2_O production in fungal and bacterial treatments (for both ^15N^BW and ^15N^Sed), and 2) N_2_O reduction by bacterial denitrifiers occurs in the ^15N^Control and ^15N^Bacterial treatments and must be added back to production estimates to correct for fractional N_2_O loss due to reduction to N_2_. These corrections ensured accurate quantification of both N_2_O production and consumption in the absence of acetylene (C_2_H_2_), which was not used in ^15^N incubations to block N_2_O reduction (Table S1).

### Natural-abundance isotope incubation experiments

NA-incubations were performed to determine the isotopic composition (δ^15^N, δ^18^O, SP) of newly produced N_2_O over a 6-day period. After the pre-incubation, 145 mL of treated unfiltered bottom water was transferred into acid-washed, muffled 160 mL serum bottles, and crimp-sealed with grey butyl rubber septa and aluminium caps. NA-sediment incubations were prepared in the same way as described for the ^15^N-sediment incubations. Eight treatments were prepared in quadruplicate: 1) ^NA^BW-Control, 2) ^NA^BW-Bacterial, 3) ^NA^BW-Fungal, 4) ^NA^BW-Chemo, 5) ^NA^Sed-Bacterial, 6) ^NA^Sed-Chemo, 7) ^NA^Sed-Fungal, 8) ^NA^Sed-Chemo. Again, a filtered bottom-water incubation without Fe(II) was included as an abiotic negative control.

To inhibit N_2_O reduction to N_2_ in a subset of samples, 1.5 ml of 100% acetylene gas (Carbagas) was injected into duplicate vials per treatment, reaching 10% v/v in the headspace (+C_2_H_2_). In the other subset of duplicate vials, no C_2_H_2_ was added to allow the potential occurrence of N_2_O reduction (Table S2).

After 30 min of He-purging, NA KNO_3_^-^ (δ^15^N = -29.6‰) was added to 10 µM final concentration. All samples were then incubated at 7 °C in the dark for 6 days. After incubation, duplicate samples per treatment were fixed by adding 3 ml of 10 M NaOH to 160 mL bottom-water bottles and 200 µL of 10 M NaOH to 20 mL sediment vials. In cases where N_2_O accumulation exceeded 15 nmol per vial, samples were split into two separate vials to allow analysis of technical replicates for ^NA^BW and ^NA^Sed incubations.

### 2.4 Rate calculations

Rates of N_2_O production (R_N2O_ in nM N/d) were calculated from the linear increase in mass 45 and 46 isotopologues of N_2_O over time in ^15^N incubation experiments.^42^ The total N_2_O production rate (R) was calculated according to Eq. 1.^43^

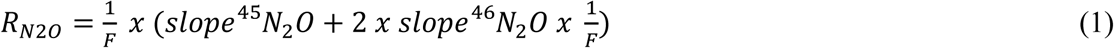

where F refers to the fraction of ^15^N in the NO_3_^-^ substrate pool (𝐹𝐹 = ^15^𝑁𝑁/ (^15^𝑁𝑁 + ^14^𝑁𝑁)), which is assumed to be constant over the duration of the incubation. The probability of ^46^N_2_O production is proportional to 1/F^2^, therefore the term includes an extra factor of 1/F for ^46^N_2_O relative to ^45^N_2_O production. F was 0.48 in the ^15N^BW-incubations and 0.78 in ^15N^Sed-incubations. Rates were considered significant at p < 0.05.

N_2_O reduction rates (R_N2O_red_ in nM N/d) were calculated from the increase in excess ^30^N_2_ over time, using the same time points as for N_2_O production rate determination (Eq. 2).^44^ No ^29^N_2_ formation was observed. The reduction rated was calculated as:

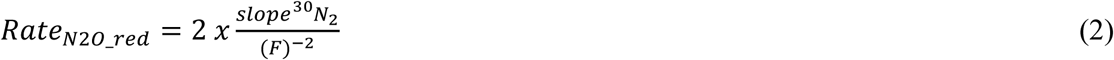

### 2.5 Modelling the relative contributions of denitrification pathways using an isotope mass balance approach

To quantify the contributions of bacterial denitrification, fungal denitrification and chemo-denitrification to N_2_O production and reduction, we applied the Fractionation and Mixing Evaluation (FRAME) model. FRAME is a Bayesian isotope mixing model with a user-friendly graphical interface (malewick.github.io/frame), recently developed by Lewicki et al^45^. The model enables simultaneous N_2_O source apportionment (and associated probability distributions) and estimation of isotope fractionation due to N_2_O reduction, using a Markov-Chain Monte Carlo approach.^45^ The model has been successfully applied in recent N_2_O source apportionment studies.^46–48^ Here, FRAME was applied to NA-control incubations of both bottom water and sediment (i.e. treatments permissive to all denitrification pathways), and an in situ bottom-water sample taken at 92.5 m depth in August.

The model used prior knowledge on isotope endmember values (SP, δ^15^N, δ^18^O) from the literature for bacterial, fungal, and chemo-denitrification as potential N_2_O sources (Table S3 and Supplemental Material). Nitrification was excluded as a N_2_O source, as all incubations and in situ water samples were anoxic.

To test the model’s ability to identify N_2_O reduction, FRAME was applied to incubation samples both with and without acetylene (C_2_H_2_). While C_2_H_2_ inhibits N_2_O reduction it may not fully suppress it. ^49–51^ Including C_2_H_2_-amended samples in the modelling efforts allowed us to estimate how much residual unreacted N_2_O the model would estimate. Notably, model outputs generally exhibited less uncertainty for incubation samples with C_2_H_2_, likely due to a cleaner N_2_O isotopic source signal (less overprinting by N_2_O reduction; Fig. S6a, d). Therefore, in the results section, we focus on results from C_2_H_2_-amended incubations. Model outputs from non-C_2_H_2_ incubations are provided in the Supplementary Material (Fig. S4a, d, Fig. S7a, f).

The model can run in 2D-isotopic space (SP and δ^18^O) or 3D (SP, δ^18^O and δ^15^N^bulk^). In this study, 3D modelling provided the most conclusive and robust source-separation constraints (see Supplementary Material for details).

## 3 RESULTS

### 3.1 Vertical distribution of nitrogen species and isotopic signatures in the South Basin

During the sampling months of June and August, water column stratification was evident, marked by a steep thermocline and oxycline (Fig. 1a, e). Surface-water temperatures ranged from 25-26 °C and decreased to 7 °C at depths of 25 m and below. The oxic-anoxic interface, defined here as the depth where O_2_ concentrations dropped below 2 µM, shifted upward from 85 m in June to 76 m in August (Fig. 1a, 1e).

**Figure 1.**
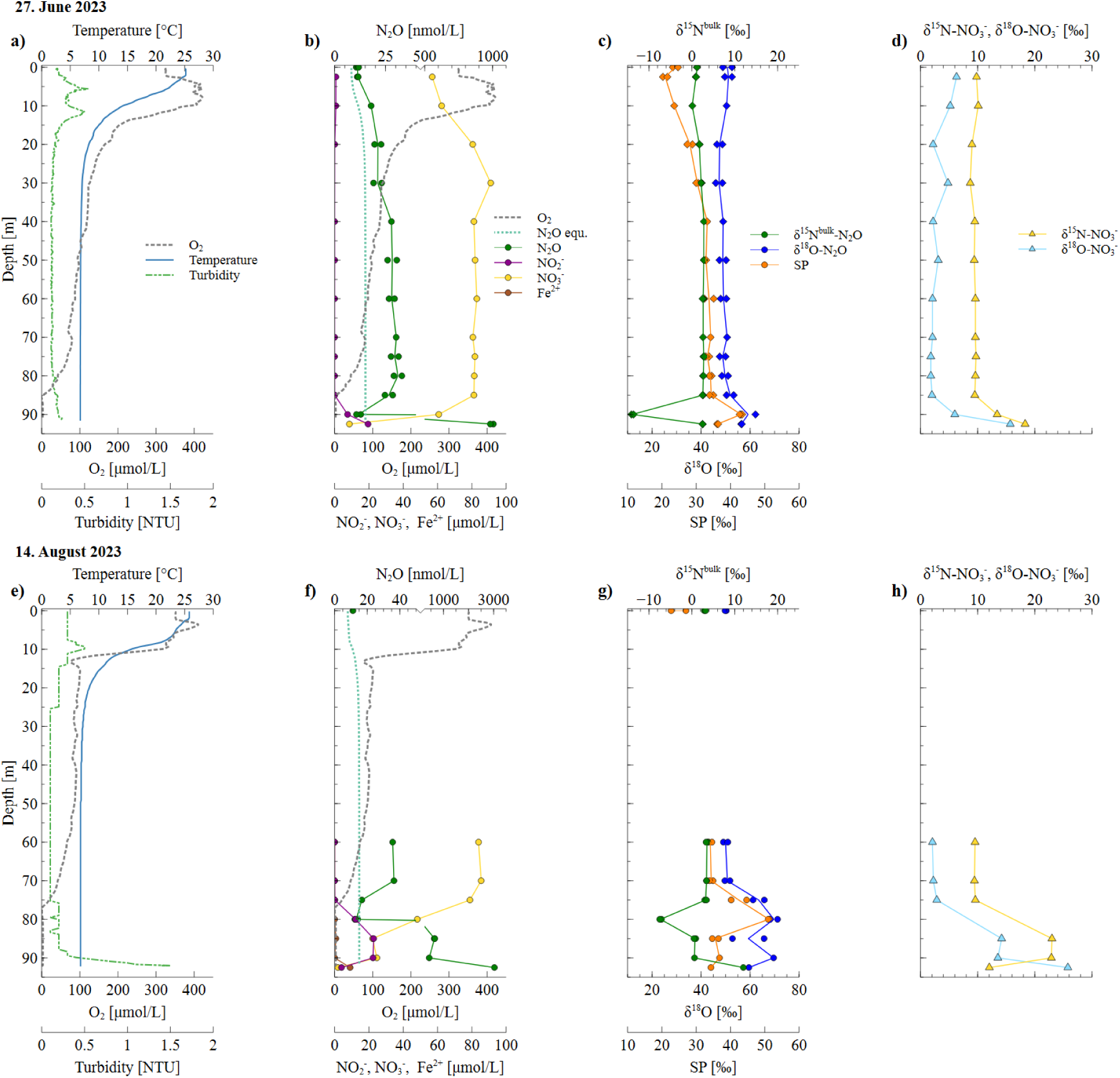
Depth profiles of physical and hydrochemical parameters in the Lake Lugano South Basin in June 2023 (upper panels) and August 2023 (lower panels). Shown are temperature, dissolved oxygen concentrations, and turbidity (**a, e**); concentrations of NO_2_^-^, NO_3_^-^, N_2_O and Fe(II), as well as calculated N_2_O concentrations for atmospheric equilibrium conditions, (**b, f**); isotopic values for N_2_O (**c, g**); and isotopic values for NO_3_^-^ (**d, h**).

Concentrations of N_2_O in surface waters reached 11 nM, slightly exceeding the atmospheric equilibrium levels (8 nM) (Fig. 1b). Concentrations increased to 34 nM at the onset of the oxycline (defined here as the water-column layer where O_2_ decreases from 40 µM to 2 µM; i.e. 80-85m in June and 70-76m in August; Fig. 1a, e), with a δ^15^N^bulk^-N_2_O value of approximately 3‰ (Fig. 1c, g). SP increased from 24‰ at the surface to 34 ‰ at the start of the oxycline in both months. δ^18^O-N_2_O mirrored SP, with values around 51‰ in the upper water column, rising to 60‰ at greater depths (Fig. 1c, g).

Within the oxycline N_2_O concentrations decreased, reaching a minimum of 12 nM in June and 13 nM in August just below the oxic-anoxic interface (Fig. 1b, f). At these depths, δ^15^N^bulk^-N_2_O values were lowest (June: -14‰ and August: -7‰, Fig. 1c, g), while SP (June: 43‰, August: 51‰) and δ^18^O (June: 60‰ August: 70‰) reached their highest values. These trends intensified, and shifted upward in August, corresponding to the upward migration of the oxic- anoxic interface.

Toward the sediment, N_2_O concentrations sharply increased, reaching 920 nM in June and 3032 nM in August in the near-bottom-water (Fig. 1b, f), where increased turbidity indicates the development of a benthic nepheloid layer^33^ (Fig. 1a, e). In August, this N_2_O accumulation was accompanied by an increase in δ^15^N^bulk^-N_2_O to 12‰, and decreases in SP and δ^18^O-N_2_O to 34‰ and 60‰, respectively (Fig. 1g).

Nitrate concentrations declined throughout the anoxic water column toward the sediment, while δ^15^N-NO_3_^-^ and δ^18^O-NO_3_^-^ values increased (Fig. 1d, h). Nitrite concentrations were below the detection limit in the oxic water column but reached a maximum of 20 µM in the lower anoxic water layer. Similarly, Fe(II) was only detected in the anoxic bottom water, with concentrations reaching 9 µM (Fig. 1f).

### 3.2 N_2_O production and N_2_O reduction rates from ^15^N-incubations

In bottom-water (BW) incubations, the highest N_2_O production rates were observed in the control treatment followed, by the bacterial treatment, which accounted for 58 ± 15% of total production (Table 1). Chemo and fungal treatments showed similar N_2_O production, representing 18 ± 2% and 24 ± 9% of the total N_2_O production rate. No N_2_O production was observed in the abiotic control (data not shown). Within error margins, the sum of chemo, fungal and bacterial-derived N_2_O production aligned with the rates observed in the control treatment, supporting the validity of the selective inhibition approach.

**Table 1.**
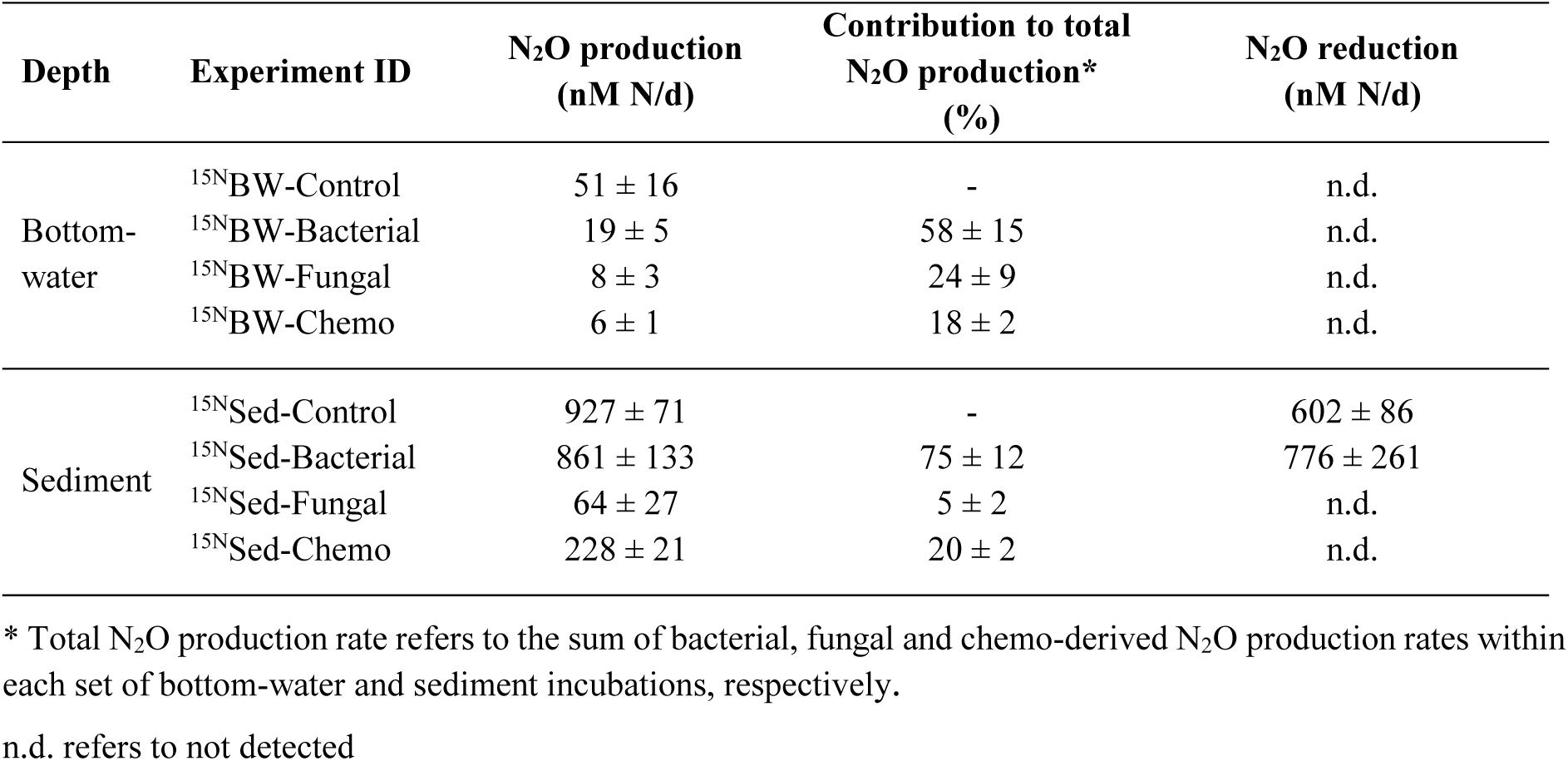
Rates of N_2_O production and reduction from ^15^N-incubation experiments. All rates are based on triplicate incubations and were significant (p <0.05).

| Depth | Experiment ID | N<sub>2</sub>O production<br>(nM N/d) | Contribution to total<br>N<sub>2</sub>O production*<br>(%) | N<sub>2</sub>O reduction<br>(nM N/d) |
| --- | --- | --- | --- | --- |
| Bottom-water | <sup>15</sup>NBW-Control | 51 ± 16 | - | n.d. |
|  | <sup>15</sup>NBW-Bacterial | 19 ± 5 | 58 ± 15 | n.d. |
|  | <sup>15</sup>NBW-Fungal | 8 ± 3 | 24 ± 9 | n.d. |
|  | <sup>15</sup>NBW-Chemo | 6 ± 1 | 18 ± 2 | n.d. |
| Sediment | <sup>15</sup>Nsed-Control | 927 ± 71 | - | 602 ± 86 |
|  | <sup>15</sup>Nsed-Bacterial | 861 ± 133 | 75 ± 12 | 776 ± 261 |
|  | <sup>15</sup>Nsed-Fungal | 64 ± 27 | 5 ± 2 | n.d. |
|  | <sup>15</sup>Nsed-Chemo | 228 ± 21 | 20 ± 2 | n.d. |

\* Total N<sub>2</sub>O production rate refers to the sum of bacterial, fungal and chemo-derived N<sub>2</sub>O production rates within each set of bottom-water and sediment incubations, respectively.
n.d. refers to not detected

Sediment incubations yielded substantially higher volumetric N_2_O production rates overall (Table 1). The control and bacterial treatments displayed the highest N_2_O production rates with the bacterial contribution, representing 75 ± 12% of total production. Chemo- and fungal denitrification showed significantly lower rates, accounting for 20 ± 2% and 6 ± 2% of the total N_2_O production rate, respectively.

N_2_O reduction was detected only in sediment incubations, specifically in the control and bacterial treatment (Table 1). No N_2_O reduction activity was observed in the fungal and chemo treatments.

### 3.3 Natural-abundance incubations – δ^18^O and site preference of accumulated N_2_O

After six days of incubation, N_2_O accumulation was much higher in sediment incubations with inhibited N_2_O reduction (+C_2_H_2_; up to 20’000 nM N_2_O) compared to BW incubations (+C_2_H_2_; up to 2’000 nM N_2_O) (Fig. S1). In the BW+C_2_H_2_ treatments, SP and δ^18^O values were similar across all inhibitor conditions (Fig. 2a, Table S5). The bacterial+C_2_H_2_ treatment yielded values within the expected range for bacterial denitrification. Unexpectedly, the chemo+C_2_H_2_ and fungal+C_2_H_2_ treatments produced comparable isotopic signatures (SP -1.8 ± 0.7‰ and 0.0 ± 0.1‰; δ^18^O 20.4 ± 0.7‰ and 22.1 ± 0.2‰, respectively). The control+C_2_H_2_ treatment showed slightly lower SP (-2.5 ± 1.2‰) and δ^18^O (17.0 ± 0.2‰) values.

**Figure 2.**
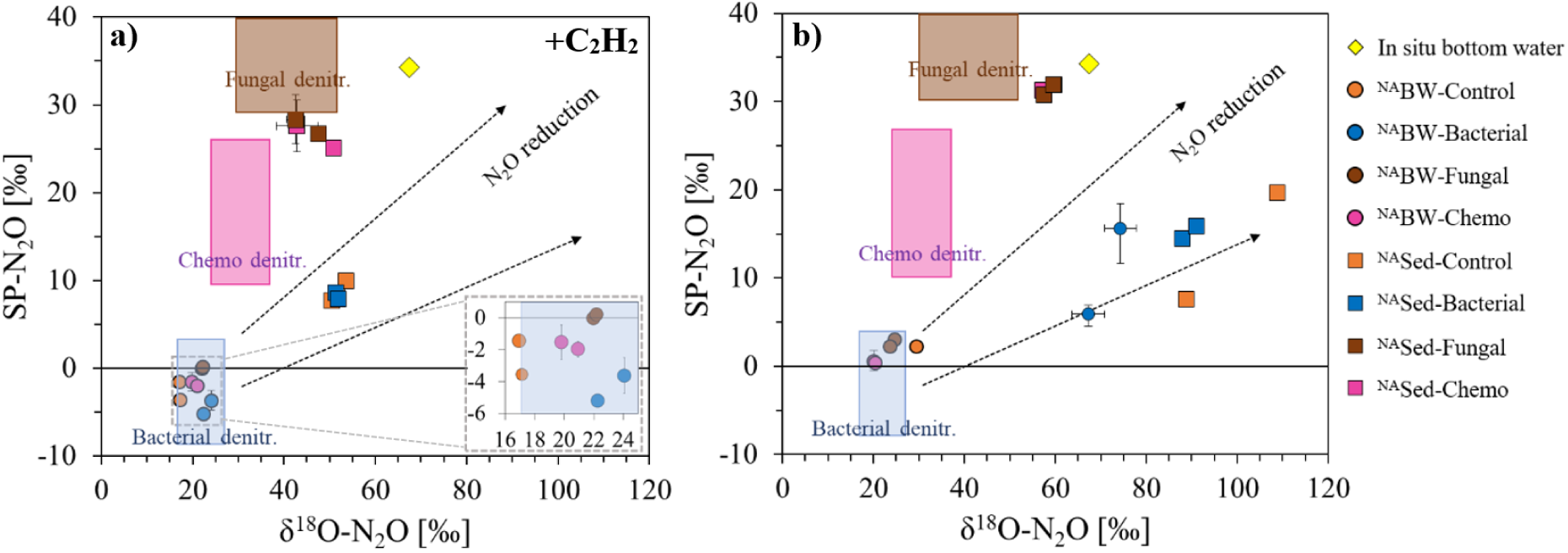
N_2_O site preference (SP) plotted against δ^18^O of N_2_O in situ bottom-water (92.5 m) and for the various treatments after six days of incubation: **a)** with C_2_H_2_ addition to inhibit N_2_O reduction, and **b)** without C_2_H_2_. The δ^18^O values are corrected for the δ^18^O of ambient water (Δδ^18^O(N_2_O – H_2_O); δ^18^O-H_2_O = -7‰^52^). Black dashed lines with arrows indicate the expected isotopic enrichment from bacterial N_2_O reduction of εSP/ε^18^O slopes ranging from of 0.23 to 0.45.^53^ The abiotic control (filtered bottom-water without Fe(II) addition) is excluded, as no N_2_O accumulation was observed. Standard errors are shown for samples where technical replicates were analysed.

In BW incubations without C_2_H_2_ (i.e., allowing N_2_O reduction), only the bacterial treatment exhibited a significant shift in the N_2_O isotopic signature, with SP increasing by 6.4‰ and δ^18^O by 47.6‰ (Fig. 2b). Other treatments exhibited isotopic signatures similar to their +C_2_H_2_ counterparts. No N_2_O accumulation occurred in the abiotic control (filtered bottom-water without Fe(II)).

In sediment incubations with C_2_H_2_, the bacterial and control treatments showed low SP values (8.2 ± 0.6‰ and 8.5 ± 1.4‰) and elevated δ^18^O values (51.6 ± 1.2‰ and 51.4 ± 1.9‰), consistent with N₂O production via bacterial denitrification. In contrast, the chemo and fungal treatments exhibited higher SP values (26.8 ± 2.5‰ and 27.5 ± 1.9‰) and slightly lower δ^18^O (45.5 ± 5.6‰ and 45.0 ± 3.2‰) (Fig. 2a), suggesting distinct production pathways from the bacterial treatment.

Without C_2_H_2_, the isotopic composition of N_2_O shifted toward higher SP and δ^18^O values in all sediment treatments (Fig. 2b). The SP and δ^18^O values of the chemo and fungal treatments showed relatively modest increases with respect to the corresponding +C_2_H_2_ treatments (∼3‰ for SP and ∼14‰ for d^18^O), while the control and bacterial treatments exhibited larger shifts (∼6‰ for SP and ∼42‰ for δ^18^O).

### 3.4 N_2_O source contributions determined by FRAME model

N_2_O source partitioning using the FRAME model, based on three isotopic parameters (SP, δ^18^O and δ^15^N^bulk^), aligned well with the results from the ^15^N-incubation experiments (Table 1 and 2). The model identified incomplete bacterial denitrification as the dominant N_2_O production pathway in both control treatments (Table 2). Smaller contributions were attributed to chemo- and fungal denitrification both in the bottom-water and sediment (Table 2).

**Table 2.** Contribution of different denitrification pathways to N_2_O accumulation in ^NA^BW/Sed-Control+C_2_H_2_ treatments and in situ bottom-water, as estimated by the FRAME model.

|  | Measured |  |  | Mass balance results |  |  |
| --- | --- | --- | --- | --- | --- | --- |
| | SP<br>[‰] | $\delta^{18}\text{O}$<br>[‰] | $\delta^{15}\text{N}^{\text{bulk}}$<br>[‰] | Bacterial<br>denitr.<br>(%) | Fungal<br>denitr.<br>(%) | Chemo<br>denitr.<br>(%) |
| <sup>NA</sup>BW-control+C<sub>2</sub>H<sub>2</sub> | -2.5 ± 1.2 | 17.0 ± 0.2 | -13.5 ± 1.0 | 88 ± 21 | 5 ± 14 | 6 ± 11 |
| <sup>NA</sup>Sed-control+C<sub>2</sub>H<sub>2</sub> | 8.5 ± 1.4 | 51.4 ± 1.9 | 16.0 ± 0.4 | 79 ± 9 | 7 ± 5 | 15 ± 10 |
| In situ bottom-water | 34.3 | 67.5 | 0.04 | 32 ± 23 | 39 ± 25 | 28 ± 24 |

We also assessed N₂O accumulation in bottom-water in situ (August sampling). FRAME- model estimates indicated similar contributions from all three pathways, though with high associated uncertainties due to model limitations (discussed in section 4.2) (Table 2).

## 4 DISCUSSION

### 4.1 N_2_O accumulation in bottom-waters originates mostly from sediments

Extremely high N_2_O accumulation (up to 3 µM) was observed in the anoxic bottom-waters of the SB, where high concentrations have been measured previously during summer stratification (up to 900 nM^31^). These N_2_O levels far exceed peak N_2_O concentrations typically observed in other lacustrine^18,54,55^ or marine systems.^14,56,57^

Sediment incubation experiments revealed that volumetric N_2_O production rates and accumulation over six days were more than an order of magnitude higher than those in bottom-water incubations (Fig. S1). This strongly suggests that most of the N_2_O in the bottom-waters originates from the sediments rather than being produced *in situ* within the water column. It should be noted that NO_3_^-^ in the in situ sediments is depleted within the first few millimetres. Hence, the sediment volume capable of sustaining reductive N_2_O production via denitrification is much smaller compared to the anoxic water volume. Nonetheless, high benthic N_2_O fluxes, around 1115 nmol N_2_O h^-1^ m^-2^, reported in previous research at the same site during summer stratification, support a sedimentary origin for the N_2_O accumulating in the bottom water.^32^ Moreover, benthic N_2_O fluxes were found to increase as stratification progresses (up to 2605 nmol N_2_O h^-1^ m^-2^ during late stage of the stratification period), whereas fluxes were much lower during the mixing period (140 to 293 nmol N_2_O h^-1^ m^-2^).^32^

The high benthic N_2_O fluxes over the stratification period result in an increase in bottom-water N_2_O concentrations from June (920 nM) to August (3032 nM). Despite the highly elevated N_2_O concentrations in near-bottom-waters, N_2_O remains undersaturated immediately below the oxic-anoxic interface, indicating that active, complete denitrification prevents N_2_O from evading into the atmosphere during the stratification. Thus, the anoxic waters below the oxic-anoxic interface function as a biogeochemical filter, impeding upward diffusion of N_2_O from sediments into the upper water column. Yet, at least at this time during the seasonal cycle, N_2_O reduction rates in the anoxic water layer are insufficient to fully counterbalance benthic N_2_O fluxes, leading to persistent N_2_O accumulation. At the end of the stratification period, this stored N_2_O could be transported into the upper water column during winter mixing, potentially rendering the SB at least transiently a significant source of N_2_O to the atmosphere. The fate of accumulated bottom-water N_2_O however remains to be further investigated. Nonetheless, these findings highlight the potential for lakes to act as pulsed sources of greenhouse gases, with implications for regional and global N_2_O budgets under changing stratification regimes.

### 4.2 Multi-isotope constraints on N_2_O source attribution

Source attribution for bottom-water and sediment incubations using the FRAME model showed strong agreement with results from ^15^N-incubation experiments, consistently identifying incomplete bacterial denitrification as the dominant N_2_O production pathway, followed by chemo-denitrification and fungal denitrification. While FRAME effectively distinguished among these processes in the controlled incubations, its application to in situ bottom-water samples led to less conclusive insight. More specifically, in the in situ samples, FRAME inferred roughly equal contributions from the three reductive N_2_O production pathways, but with high uncertainties (Table 2, Fig. S8a, b). In fact, when initially only SP and δ^18^O were used as isotopic input parameters, FRAME misleadingly suggested fungal denitrification as the dominant source of N_2_O in the in situ bottom-water (Fig. S5a, b). However, results from both ^15^N- and NA-incubations clearly indicated bacterial denitrification as the primary source, with fungal denitrification playing only a minor role. This obvious discrepancy prompted the refinement of the FRAME model through the inclusion of a third parameter (δ^15^N^bulk^), which "improved" the source attribution output by reducing the overestimation of fungal denitrification due to the extension of the mixing space. However, including δ^15^N^bulk^ as a parameter led to source apportionment with large standard deviations, indicating that the model failed to converge on a probable solution. This is evident from the pronounced oscillations of the Markov chain for the in situ bottom-water sample, reflecting that maximum likelihood was not achieved (Fig. S8). The high uncertainties in the in situ sample source-attribution results expose limitations of the FRAME model, particularly when isotopic signatures fall outside the mixing space, or when N_2_O reduction isotope fractionation effects are not well constrained, and strong isotopic overprinting due to N_2_O reduction is expected (see Supplementary Material, Figs. S3–S8).

This highlights the need to consider multiple isotopic parameters and contextual information to assess and improve the reliability and accuracy of process identification and N_2_O source attribution (e.g., information about the study site can help exclude certain pathways, such as nitrification as a source of N_2_O due to anoxic conditions; or, data on substrate isotopic composition can better constrain the range of isotopic endmembers, and independent ^15^N rate measurements can be used to verify model outputs) . Nonetheless, more precise baseline data on the isotope effects associated with different N_2_O production/consumption processes are still needed. For instance, recent work suggests that the N_2_O isotope signature of bacterial denitrification may extend to higher SP values (∼10‰).^28^ Furthermore, N_2_O production via DNRA performed by bacteria belonging to the family of *Geobacteraceae* was reported to exhibit SP values >43‰.^8^ This would clearly impact future model calibrations. Despite these challenges, the use of canonical literature values for isotopic endmembers used in the FRAME model remains a valuable tool for disentangling N_2_O production, particularly when combined with targeted incubation experiments and careful site-specific validation.

### 4.3 Incomplete bacterial denitrification dominates N_2_O production in bottom water and sediments

Both ^15^N- and NA-incubation experiments identified incomplete bacterial denitrification as the dominant N_2_O production mechanism. Bacterial N_2_O production accounted for 58 ± 15% in the bottom water and 68 ± 3% in sediments, based on direct rate measurements, and 88 ± 21% and 79 ± 9%, respectively, based on the isotope mass balance approach.

In bottom-water treatments, the addition of a bacterial inhibitor (STP) was expected to suppress bacterial N_2_O production. However, the isotopic signatures (SP and δ^18^O) of the N_2_O produced remained characteristic of bacterial denitrification (Fig. 2a). This suggests incomplete inhibition of bacterial activity (i.e., in the ^NA^BW-Chemo+C_2_H_2_ and the ^NA^BW-Fungal+C_2_H_2_ treatments). Given that fungal and chemo-denitrification contributions were minimal in the ^NA^BW incubations, even low residual bacterial activity could dominate the N_2_O isotopic signal. As with many inhibitor-based approaches, full inhibition is often difficult to ensure and may be accompanied by non-target effects by the inhibitors.^58–60^ Further supporting bacterial dominance, the isotopic data for all bottom-water treatments (including the bacteria-inhibition treatments targeting fungal and chemo-denitrification) plotted within the bacterial endmember box in the SP-vs.-δ^15^N^bulk^ space (Fig. S2a, b). Nevertheless, a closer examination reveals that chemo and fungal treatments exhibited slightly elevated SP values compared to the bacterial control (zoom-in of Fig. 2a), suggesting limited but detectable activity of these alternative N_2_O production processes.

In sediment incubations, the control and bacterial treatment had overlapping SP and δ^18^O signatures, again pointing to bacterial denitrification as the primary N_2_O source (Fig. 2a). Notably, SP and δ^18^O values in the bacterial treatment were higher than typically reported for bacterial denitrification. Several factors may explain this: Firstly, the range of SP values for bacterial denitrification was recently extended following the identification of a NO-detoxifying enzyme (fhp), widely present in marine and terrestrial denitrifying bacteria, which produces N_2_O with a SP of 10‰.^28^ This suggests that bacterial denitrification can produce N_2_O with SP values above the traditionally accepted range of -8 to 4‰. Additionally, DNRA has been shown to produce N_2_O with SP values exceeding +43‰.^8^ The contribution of DNRA to total benthic NO_3_^-^ reduction in Lake Lugano’s South Basin was reported to be less than 12% in a study by Wenk et al.^38^. However more recent research indicated that DNRA may contribute up to ∼50% at the same study site.^39^ Despite this, a high contribution of N_2_O production from DNRA is unlikely in our study, as the SP value of ∼8‰ in the control and bacterial treatments (i.e., the only treatments that permitted the occurrence of DNRA) was much lower than the SP values typically associated with DNRA. This suggests that DNRA played only a minor role, and that bacterial denitrification was the dominant process. Nonetheless, a relatively small contribution of DNRA could explain the shift in SP values to ∼8‰. in the bacterial and control treatments Elevated SP and δ^18^O values may also result from incomplete inhibition of either N_2_O reduction to N_2_ or fungal denitrification. Indeed, the effectiveness of C_2_H_2_ as an inhibitor of N_2_O reduction is known to vary.^49–51^ Lastly, in the bacterial treatment, chemo-denitrification was not specifically inhibited, and its contribution could have shifted the isotopic signatures towards SP and δ^18^O values that are higher than typically associated with bacterial denitrification.

Despite these complexities, the consistently high benthic N_2_O reduction rates observed in both control and bacterial treatments further support bacterial denitrification as the dominant N_2_O source in the sediment and bottom water Lake Lugano’s South Basin.

### 4.4 Isotopic signatures of N_2_O in the bottom-waters reflect N_2_O reduction in sediments

The elevated SP values of N_2_O observed in the bottom water cannot be explained by N_2_O production via incomplete bacterial denitrification in the water column alone (i.e., they are uncharacteristic for N_2_O production by bacterial denitrification, which typically yields lower isotopic signatures). Instead, these "enriched values" are best explained by microbial N_2_O reduction occurring within the sediments, where the majority of the bottom-water N_2_O originates from, and where we demonstrated N_2_O reduction to occur concurrently (Table 1). While N_2_O reduction rates in bottom-water incubations were below detection limit (Table 1), sediment incubations clearly demonstrated the effect of N_2_O reduction on the N_2_O isotopic composition. In ^NA^Sed incubations without C_2_H_2_ (i.e., a N_2_O reduction inhibitor), only the control and bacterial treatments showed a strong shift toward elevated SP and δ^18^O values (Fig. 2b), along with significantly lower N_2_O concentrations compared to C_2_H_2_-amended treatments (Fig. S1). This directly links the observed isotopic enrichment to active N_2_O reduction. As expected, no N_2_O reduction was measured in the chemo- and fungal denitrification treatments (i.e., bacteria-inhibited conditions; Table 1), consistent with the understanding that bacteria are the primary N_2_O reducers.^19,61^

Together, these results demonstrate that the enriched SP and δ^18^O values in bottom-water N_2_O are primarily driven by microbial reduction processes in the sediment, rather than reflecting distinct N_2_O production pathways alone. This highlights the importance of considering reduction dynamics when interpreting isotopic data from stratified aquatic systems.

### 4.5 Secondary but non-negligible role of chemo-denitrification and fungal denitrification

Chemo-denitrification and fungal denitrification contributed much less to N_2_O production than bacterial denitrification in the Lake Lugano southern basin, but their roles, especially that of chemo-denitrification, were nonetheless substantial. Together, they represent non-canonical but significant pathways of N_2_O formation, which are often overlooked in biogeochemical models and source attribution studies.

Chemo-denitrification accounted for approximately 20 ± 2% in sediment and 18 ± 2% in bottom water, based on ^15^N tracer experiments. Even when considering the possibility of incomplete inhibition in some experiments, these values indicate a non-negligible contribution, particularly for a pathway that is not traditionally emphasized in aquatic N_2_O budgets. Previous studies have shown that chemo-denitrification can contribute 13 to 28% of total N_2_O production in estuarine sediments,^16^ and up to 70% under elevated NO_3_^-^ conditions in coastal sediments.^17^

The activity of chemo-denitrification has been reported to be enhanced by the presence of Fe(II)-bearing minerals.^62,63^ Under natural, pH-neutral conditions, the majority of aqueous Fe(II) binds to mineral surfaces or ligands, forming Fe(II)-bound phases that act both as Fe^2+^ sources as well as a reactive surfaces for NO_2_^-^ reduction.^62^ In the Lake Lugano South Basin, dissolved Fe^2+^ concentrations were higher than those of particulate Fe(II), and Fe concentrations increased significantly in the sediment.^41^ This likely explains the higher chemo- denitrification rates observed in sediments compared to the water column (Table 1).

Fungal denitrification contributed modestly to N_2_O production in the sediments, with both NA- and ^15^N-incubation approaches estimating a similar contribution of ∼6%. However, results for the bottom-water incubations were inconsistent. The ^15^N approach resulted in a fungal N_2_O production rate of 5 ± 2 nM N/d, contributing 24 ± 9% of total N₂O production, while the isotope mass balance approach suggested a much smaller contribution (5 ± 14%). This discrepancy may stem from the overall low N₂O production in bottom-waters, where even small absolute contributions/differences can appear large in relative terms, and can therefore result in larger errors. Moreover, despite the use of a bacterial inhibitor in the ^15^N-label incubations, the N_2_O produced in the fungal treatment may still reflect residual bacterial activity (i.e., incomplete inhibition). This hypothesis is supported by bottom-water NA- incubations, where, as mentioned above, SP values in both chemo- and fungal denitrification treatments were consistent with bacterial denitrification (Fig. 2). A slight shift toward higher SP values in the fungal treatment compared to the bacterial treatment may indicate a somewhat greater relative (but still minor) contribution to total N_2_O production from fungal denitrifiers. Further investigations are needed to fully understand and confirm the role of fungi in N_2_O cycling in Lake Lugano, and other aquatic systems.

Overall, while fungal denitrification appears to play a minor role in Lake Lugano, and is difficult to isolate with certainty, both chemo-denitrification and fungal pathways contribute to N_2_O production in the lake’s southern basin. These findings highlight the importance of accounting for non-canonical reductive N_2_O sources beyond bacterial denitrification, both in Lake Lugano and in other lake systems. Their relevance may increase under elevated NO_3_^-^ conditions or shifting redox regimes,^16,17^ making them increasingly significant components of N_2_O cycling in eutrophic aquatic environments.

## ASSOCIATED CONTENT

### Supporting information

The Supporting information is available free of charge on the ACS Publications website at DOI:

Literature values for N_2_O production of different denitrification pathways (Table S3) and for N_2_O reduction (Table S4), Isotopic values for NA-incubation experiments (Table S5), N_2_O concentrations in NA-incubation experiments (Fig. S1), dual isotope plot (SP vs δ^15^N) for NA-incubation samples, FRAME model approach description and outputs (Fig. S3-S8).

### Author Contributions

T.E. and C.F. designed research and planned experiments; T.E. performed experiments and analysed data; all authors discussed the results; T.E. wrote the paper with contributions from all authors.

### Funding Sources

This work was funded by the Swiss National Science Foundation (Grant number 201027).

### Notes

The authors declare no competing financial interest

## ACKNOWLEDGMENTS

We thank Camilla Capelli and Fabio Lepori from the University of Applied Sciences and Arts of Southern Switzerland (SUPSI) for support during sampling. We thank Thomas Kuhn and Franz Conen for help with N_2_O concentration and isotope analysis.

