## Supplementary Information for "Non-canonical reductive nitrous oxide production pathways in a seasonally stratified lake basin"

### Content of the file:

Supporting Tables

Table S1: Overview of $^{15}N$-labelled incubation experiments

Table S2: Overview of natural-abundance incubation experiments

Table S3: Literature values for $N_2O$ production of different denitrification pathways

Table S4: Literature values for $N_2O$ reduction

Table S5: Isotopic values for NA-incubation experiments

Supporting Figures

Figure S1: $N_2O$ concentrations in NA-incubation experiments

Figure S2: Dual isotope plot (SP vs $\delta^{15}N$) for NA-incubation samples

FRAME model approach description and outputs (Figure S3-S8)

Reference list

**Table S1.** Overview of $^{15}$N-labelled $NO_3^-$ incubation experiments performed to quantify $N_2O$ production via bacterial-, fungal- or chemo-denitrification, respectively, in bottom water and sediment from Lake Lugano. Treatments were distinguished by the use of selective microbial inhibitors. Streptomycin (STP) and Cycloheximide (CYH) were used to suppress bacterial and fungal denitrification, respectively.

| Set | Depth | ID | $C_2H_2$ addition | Active denitrification pathway | Inhibitor addition | Fe(II) addition | Correction of $N_2O$ production rate for | Substrate | Analysed for |
|---|---|---|---|---|---|---|---|---|---|
| $^{15}$N | Bottom-water (BW) | $^{15N}$BW-Control | no | Control (chemo-, fungal, bacterial denitr.) | no inhibitor | 2 µM | - | 10 µM of $^{15}$N-labelled $KNO_3$ | Production of $^{45}N_2O$ and $^{46}N_2O$; Production of $^{30}N_2$ |
| | | $^{15N}$BW-Bacterial | | Bacterial denitr. (+ chemo-denitr.) | 5.33 mM CYH | 2 µM | corrected for $N_2O$ produced through chemo-denitr; | | |
| | | $^{15N}$BW-Fungal | | Fungal denitr. (+ chemo-denitr.) | 1.29 mM STP | 2 µM | corrected for $N_2O$ produced through chemo-denitr. | | |
| | | $^{15N}$BW-Chemo | | Chemo-denitr. | 5.33 mM CYH + 1.29 mM STP | 2 µM | - | | |
| $^{15}$N | Sediment (Sed) | $^{15N}$Sed-Control | | Control (chemo-, fungal, bacterial denitr.) | no inhibitor | no* | corrected for $N_2O$ loss through $N_2O$ reduction | 10 µM of $^{15}$N-labelled $KNO_3$ | Production of $^{45}N_2O$ and $^{46}N_2O$; Production of $^{30}N_2$ |
| | | $^{15N}$Sed-Bacterial | | Bacterial denitr. (+ chemo-denitr.) | 5.33 mM CYH | no* | corrected for $N_2O$ produced through chemo-denitr; corrected for $N_2O$ loss through $N_2O$ reduction | | |
| | | $^{15N}$Sed-Fungal | | Fungal denitr. (+ chemo-denitr.) | 1.29 mM STP | no* | corrected for $N_2O$ produced through chemo-denitr | | |
| | | $^{15N}$Sed-Chemo | | Chemo-denitr. | 5.33 mM CYH + 1.29 mM STP | no* | - | | |

* no Fe(II) addition is necessary due to the large solid-phase Fe(II) pool and continuous $Fe^{2+}$ production in these sediment.[1]

**Table S2**. Overview of natural-abundance (NA)-incubation experiments performed to determine the isotopic signature of accumulated $N_2O$ via bacterial-, fungal- or chemo-denitrification, respectively, in bottom water and sediment from Lake Lugano. Treatments were distinguished by the use of selective microbial inhibitors. Streptomycin (STP) and Cycloheximide (CYH) were used to suppress bacterial and fungal denitrification, respectively.

| Set | Depth | ID | $C_2H_2$ addition | Active denitrification pathway | Inhibitor addition | Fe(II) addition | Substrate | Analysed for |
|---|---|---|---|---|---|---|---|---|
| Natural-abundance (NA) | Bottom-water (BW) | $^{NA}$BW-Control+$C_2H_2$ † | yes | Control (chemo-, fungal, bacterial denitr.) | no inhibitor | 2 µM | 10 µM of $KNO_3$ | $\delta^{15}N^{bulk}$, $\delta^{18}O$, SP, $N_2O$ concentration |
| | | $^{NA}$BW-Control | no | | | | | |
| | | $^{NA}$BW-Bacterial+$C_2H_2$ | yes | Bacterial denitr. (+ chemo-denitr.) | 5.33 mM CYH | 2 µM | | |
| | | $^{NA}$BW-Bacterial | no | | | | | |
| | | $^{NA}$BW-Fungal+$C_2H_2$ | yes | Fungal denitr. (+ chemo-denitr.) | 1.29 mM STP | 2 µM | | |
| | | $^{NA}$BW-Fungal | no | | | | | |
| | | $^{NA}$BW-Chemo+$C_2H_2$ | yes | Chemo-denitr. | 5.33 mM CYH + 1.29 mM STP | 2 µM | | |
| | | $^{NA}$BW-Chemo | no | | | | | |
| Natural-abundance (NA) | Sediment (Sed) | $^{NA}$Sed-Control+$C_2H_2$ † | yes | Control (chemo-, fungal, bacterial denitr.) | no inhibitor | no* | 10 µM of $KNO_3$ | $\delta^{15}N^{bulk}$, $\delta^{18}O$, SP, $N_2O$ concentration |
| | | $^{NA}$Sed-Control | no | | | | | |
| | | $^{NA}$Sed-Bacterial+$C_2H_2$ | yes | Bacterial denitr. (+ chemo-denitr.) | 5.33 mM CYH | no* | | |
| | | $^{NA}$Sed-Bacterial | no | | | | | |
| | | $^{NA}$Sed-Fungal+$C_2H_2$ | yes | Fungal denitr. (+ chemo-denitr.) | 1.29 mM STP | no* | | |
| | | $^{NA}$Sed-Fungal | no | | | | | |
| | | $^{NA}$Sed-Chemo+$C_2H_2$ | yes | Chemo-denitr. | 5.33 mM CYH + 1.29 mM STP | no* | | |
| | | $^{NA}$Sed-Chemo | no | | | | | |

† results used for mass balance approach

* no Fe(II) addition is necessary due to the large Fe(II) pool and continuous $Fe^{2+}$ production in these sediments.[1]

**Table S3**: Isotopic signature ranges from the literature for $N_2O$ production of different denitrification pathways.

| **Process** | **$\delta^{15}N^{bulk}$** | **$\delta^{18}O$** | **SP** | **Reference** |
|---|---|---|---|---|
| Bacterial denitr. | -30 to 2.5‰ | 17 to 27‰ | -7.5 to 3.7‰ | [2–6] |
| Fungal denitr. | 4 to 70‰ | 30 to 52‰ | 30 to 39‰ | [7–9] |
| Chemo-denitr. | 16 to 38‰ | 24 to 37‰ | 10 to 27‰ | [10,11] |

**Table S2**: Isotopic signature for $N_2O$ reduction taken from Yu et al.[6]

| | |
|---|---|
| $\varepsilon SP_{N2Ored}$ | -8.2 to -2.9‰, mean -5.9‰ |
| $\varepsilon^{18}O_{N2Ored}$ | -25.1 to -5.1‰, mean -15.4‰ |
| $\varepsilon^{15}N_{N2Ored}$ | -11.0 to -1.8‰, mean -7.1‰ |
| $\varepsilon SP_{N2Ored}/\ \varepsilon^{18}O_{N2Ored}$ | 0.23 to 0.45 |
| $\varepsilon SP_{N2Ored}/\ \varepsilon^{15}N_{N2Ored}$ | 0.51 to 0.96 |

**Table S4**: $N_2O$ site preference (SP), $\delta^{18}O$ and $\delta^{15}N^{bulk}$ values from NA incubation experiments. $N_2O$ $\delta^{15}N^{bulk}$ values are normalized to the $\delta^{15}N$ of the substrate, and $N_2O$ $\delta^{18}O$ values to the $\delta^{18}O$ of ambient water ($\delta^{15}N$-$NO_3^-$ = -29.6 ± 0.3‰ and $\delta^{18}O$-$H_2O$ = -7.0‰) (n = 2).

| **Depth** | **ID** | **$\delta^{15}N$-$N_2O$ [‰]** | **$\delta^{18}O$-$N_2O$ [‰]** | **SP [‰]** |
|---|---|---|---|---|
| Bottom water (BW) | $^{NA}$BW-control+$C_2H_2$ | -13.5 ± 1.0 | 17.0 ± 0.2 | -2.5 ± 1.2 |
| | $^{NA}$BW-control † | -15.4 ± 0.1 | 29.4 ± 0.5 | 2.2 ± 0.4 |
| | $^{NA}$BW-chemo+$C_2H_2$ | -7.1 ± 1.0 | 20.4 ± 0.7 | -1.8 ± 0.7 |
| | $^{NA}$BW-chemo | -7.2 ± 0.5 | 20.2 ± 0.4 | 0.6 ± 0.4 |
| | $^{NA}$BW-fungal+$C_2H_2$ | -8.0 ± 0.2 | 22.1 ± 0.2 | 0.0 ± 0.1 |
| | $^{NA}$BW-fungal | -11.3 ± 1.8 | 27.0 ± 5.6 | -0.2 ± 6.1 |
| | $^{NA}$BW-bacterial+$C_2H_2$ | -8.6 ± 2.5 | 23.2 ± 1.0 | -4.4 ± 1.1 |
| | $^{NA}$BW-bacterial | -3.9 ± 4.5 | 70.8 ± 4.8 | 10.8 ± 5.9 |
| Sediment (Sed) | $^{NA}$Sed-control+$C_2H_2$ | 3.9 ± 0.4 | 51.4 ± 1.9 | 8.5 ± 1.4 |
| | $^{NA}$Sed-control † | -9.1 ± 4.7 | 98.9 ± 14.2 | 13.6 ± 8.6 |
| | $^{NA}$Sed-chemo+$C_2H_2$ | 18.8 ± 1.0 | 45.5 ± 5.6 | 26.8 ± 2.5 |
| | $^{NA}$Sed-chemo | 16.6 ± 0.1 | 58.5 ± 1.8 | 31.6 ± 0.5 |
| | $^{NA}$Sed-fungal+$C_2H_2$ | 19.1 ± 0.2 | 45.0 ± 3.2 | 27.5 ± 1.9 |
| | $^{NA}$Sed-fungal | 16.3 ± 0.3 | 58.6 ± 1.4 | 31.3 ± 0.8 |
| | $^{NA}$Sed-bacterial+$C_2H_2$ | 4.3 ± 0.4 | 51.6 ± 1.2 | 8.2 ± 0.6 |
| | $^{NA}$Sed-bacterial | 0.7 ± 1.2 | 89.5 ± 2.2 | 15.2 ± 1.0 |

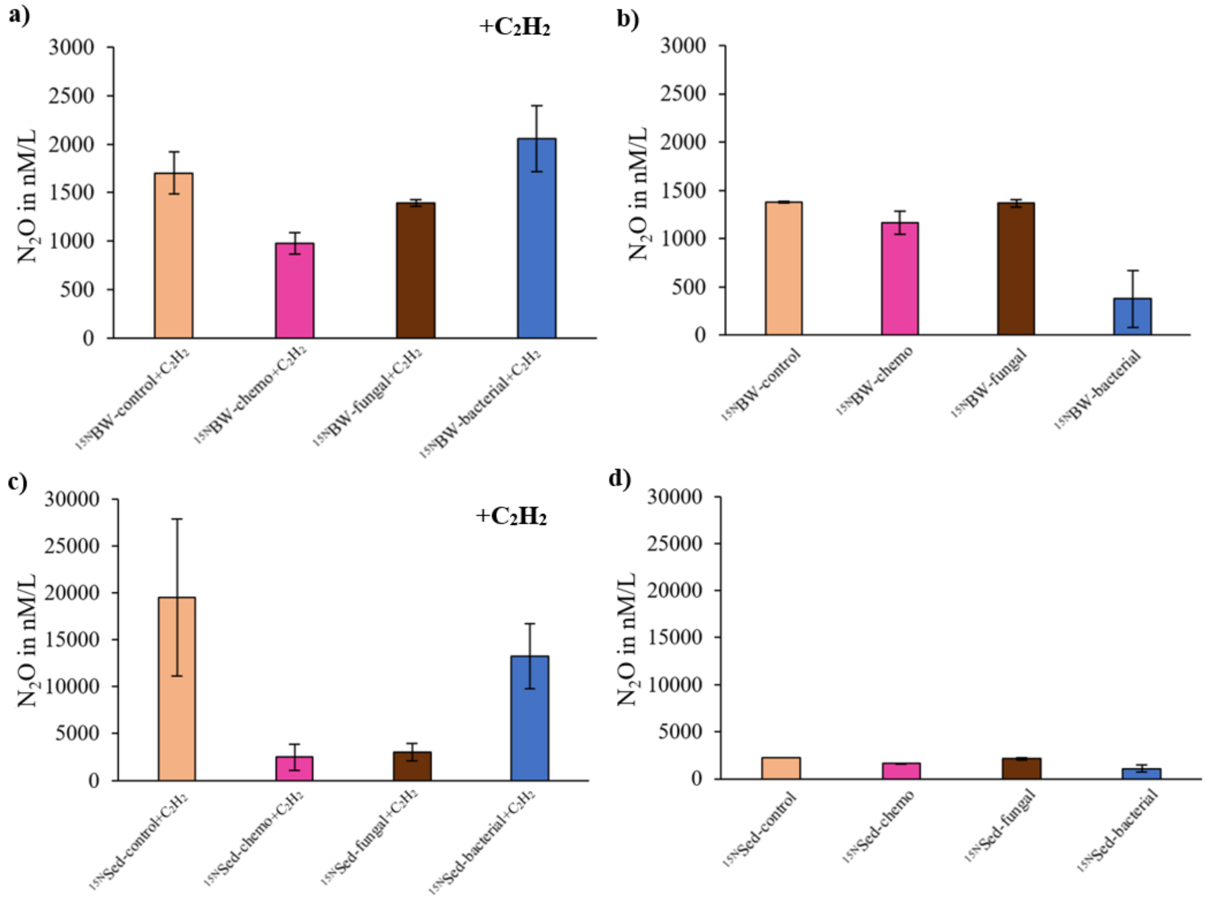


**Fig. S1**: $N_2O$ accumulation after six days of incubation in: **a**) $^{NA}$BW incubations with $C_2H_2$ and **b**) without $C_2H_2$, **c**) $^{NA}$Sed incubations with $C_2H_2$, and d) $^{NA}$Sed incubations without $C_2H_2$.

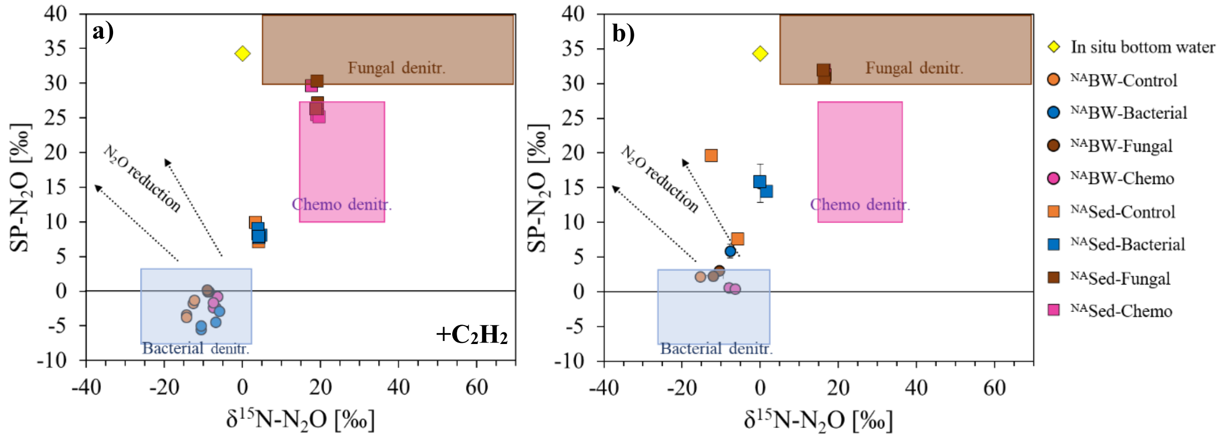


**Fig. S2**: $N_2O$ site preference (SP) and $\delta^{15}N$-values (‰) of different treatments from NA incubations with **a**) $C_2H_2$ addition and **b**) without $C_2H_2$ addition. $\delta^{15}N$-$N_2O$ is normalized to the $\delta^{15}N$ of the substrate ($\Delta\delta^{15}N(NO_3^- - N_2O)$; $\delta^{15}N$-$NO_3^-$ = -29.6 ± 0.3%). Black dashed arrows indicate the slope of the $N_2O$ reduction line (-0.51 to -0.96).[9]

### FRAME model approach description and outputs

The FRAME model was applied to control treatments and to the in situ bottom-water sample (92.5 m August), incorporating bacterial, fungal and chemo-denitrification as isotopic endmembers. Additionally, the $N_2O$ isotopic composition can be altered by the isotope fractionation induced by $N_2O$ reduction. Different approaches were used to account for this effect in incubation vs. in situ samples: For the incubation samples (closed system), a Rayleigh-type fractionation model was applied (Eq. 1):

$$\delta_{final} = \delta_{inital} + \varepsilon_{N2Ored}\ x\ ln(f_{residual}) \quad (1)$$

For the in situ samples (open-system with continuous $NO_3^-$ supply), an open-system fractionation model was used (Eq. 5):

$$\delta_{final} = \delta_{initial} - \varepsilon_{N2Ored}\ x\ (1 - f_{residual}) \quad (5)$$

In both cases, $\delta_{final}$ is the measured isotopic composition of $N_2O$ after reduction, and $\delta_{initial}$ is the isotopic composition of the initial mixture before $N_2O$ reduction. $\varepsilon_{N2Ored}$ represents the kinetic isotope effect of $N_2O$ reduction, based on literature values (Table S4), and $f_{residual}$ is the fraction of unreacted $N_2O$.

Initially, a 2D-version of the FRAME model (only SP and $\delta^{18}O$ as isotopic parameters) was tested. However, it failed to converge for the $^{NA}$BW-control+$C_2H_2$ sample (indicated by the absence of burnout, reflecting that the model did not reach maximum likelihood; Fig. S3e) and exhibited large uncertainties for the $^{NA}$Sed-control (Fig. S4b). Furthermore, the 2D output estimated a high contribution of $N_2O$ from fungal denitrification (76 ± 16%) and a substantial fraction of residual, unreacted $N_2O$ (90 ± 8%) for the in situ bottom-water sample (Fig. S5a). These results contradicted the more discrete findings from $^{15}N$-label incubation experiments, which identified bacterial denitrification as the dominant $N_2O$ production pathway.

To improve model constraints, we included $\delta^{15}N^{bulk}$ as a third parameter. This expanded the model into a 3D mixing space, which provided better source attribution for the incubation samples. The 3D approach yielded narrowly constrained probability distributions for the control samples with $C_2H_2$, clearly identifying bacterial denitrification as the dominant $N_2O$ source (Fig. S6a, b), in good agreement with findings from the $^{15}N$ labelling approach. Surprisingly, despite the addition of $C_2H_2$ to inhibit $N_2O$ reduction, a relatively low fraction of residual (i.e., unreacted) $N_2O$ was estimated for the $^{NA}$Sed-control+$C_2H_2$ (35 ± 4%, Fig. S6b), indicating that approximately 65% of $N_2O$ was consumed in this sample. Incomplete inhibition of $N_2O$ reduction in $C_2H_2$-amended samples has been previously reported.[12–14] Nonetheless, source apportionment was considered reliable due to the narrow probability distributions and the good agreement with $^{15}N$ incubation results. For the in situ bottom-water sample, however, source apportionment remained challenging, even with the 3D approach. While the 3D model no longer identified fungal denitrification as the dominant source (as the 2D model did), it instead suggested approximately equal contributions from all three production pathways, albeit with considerable uncertainty. This indicated that the 3D model was unable to identify a high-probability solution for the in situ bottom-water sample.

Given that $^{15}N$-label incubations showed fungal denitrification to be a minor pathway in the Lake Lugano southern basin, we consider the 2D output to be inaccurate for the in situ bottom-water sample. Furthermore, as the 3D approach yielded improved results for the incubation samples, we rely on the 3D results in this manuscript, despite the associated uncertainties for the in situ bottom-water sample. This illustrates both the limitations of isotope-based source apportionment modelling and, for "calibrating purposes", the importance of complementary incubation experiments, as further discussed in the main manuscript.

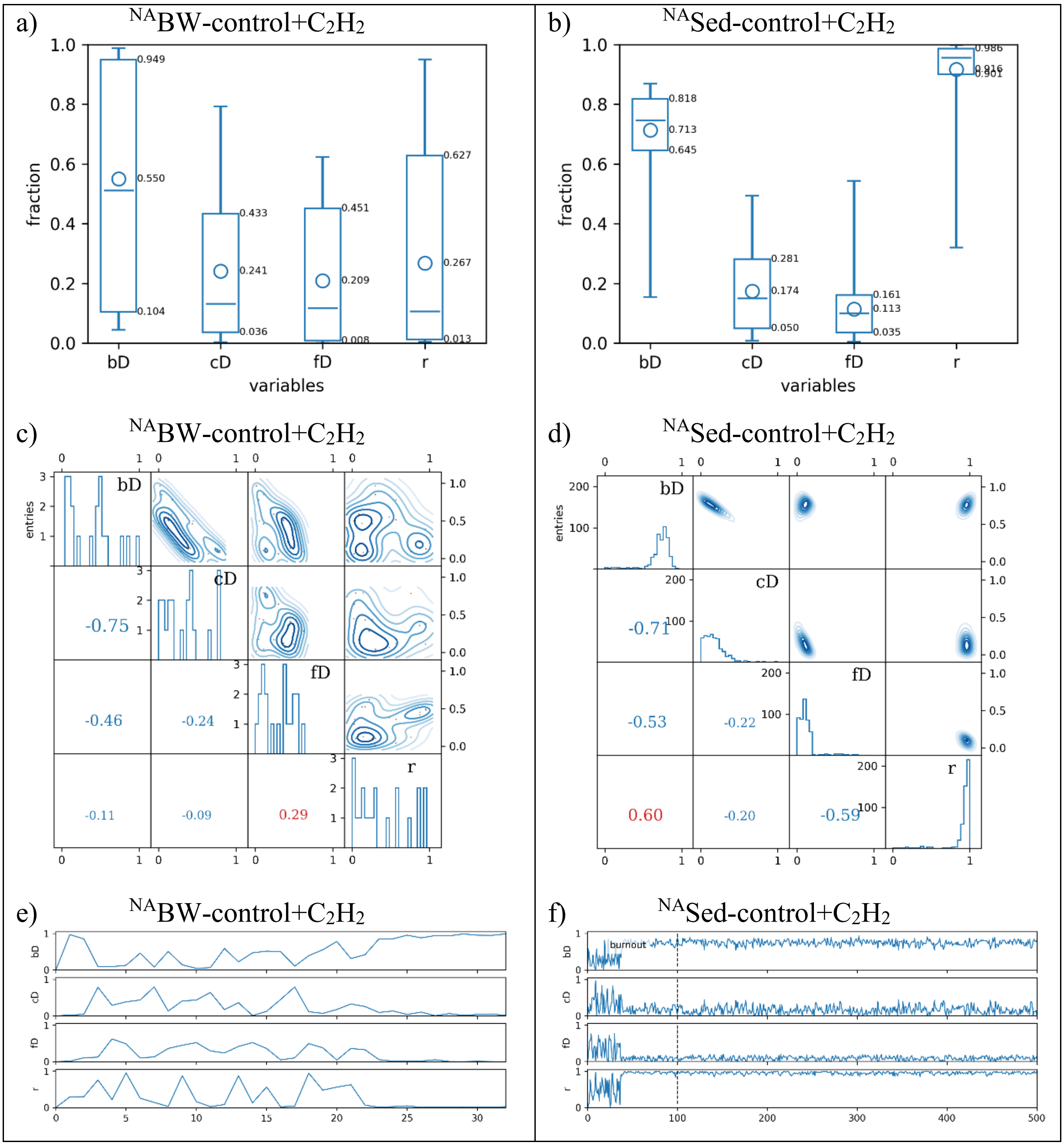


**Fig S3**: 2D FRAME model output for $^{NA}$BW-control+$C_2H_2$ and $^{NA}$Sed-control+$C_2H_2$ samples. Panels show: (**a, b**) Summary plot of distributions with key statistics (mean = circle, median = horizontal line, 68% confidence interval (CI) = box, 95% CI = whiskers); (**c, d**) Histograms showing the probability distributions of mixing fractions (diagonal), with correlation displayed as contour plots (upper right), and numerical correlation coefficients (lower left); (**e, f**) Markov chain illustrating model iteration progress, including mixing configurations and the number of iterations needed to reach stabilization. Only data after the point of stabilization (dashed vertical line) are used, and iterations before are discarded and, marked as burnout. Burnout was not reached for the $^{NA}$BW-control+$C_2H_2$ sample, whereas burnout was successfully reached for the $^{NA}$Sed-control+$C_2H_2$ sample marked by the vertical dashed line.

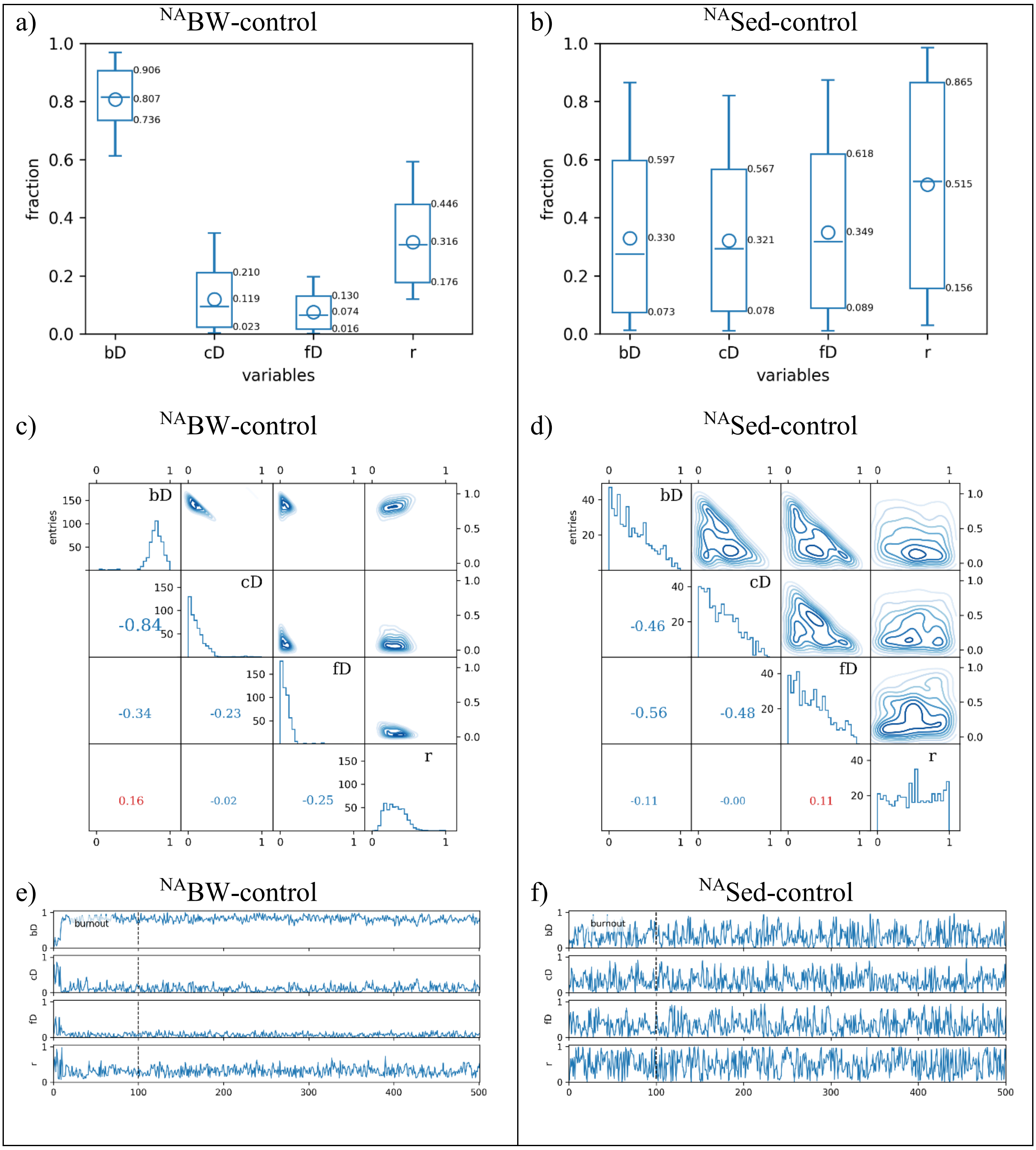


**Fig. S4**: 2D FRAME model output for $^{NA}$BW-control and $^{NA}$Sed-control samples. Panels show: (**a, b**) Summary plot of distributions with key statistics (mean = circle, median = horizontal line, 68% confidence interval (CI) = box, 95% CI = whiskers); (**c, d**) Histograms showing the probability distributions of mixing fractions (diagonal), with correlation displayed as contour plots (upper right), and numerical correlation coefficients (lower left); (**e, f**) Markov chain illustrating model iteration progress, including mixing configurations and the number of iterations needed to reach stabilization. Only data after the point of stabilization (dashed vertical line) are used, and iterations before are discarded, and marked as burnout. For $^{NA}$Sed-control, large oscillations in the Markov chain indicate that the model could not identify a stable or probable solution.

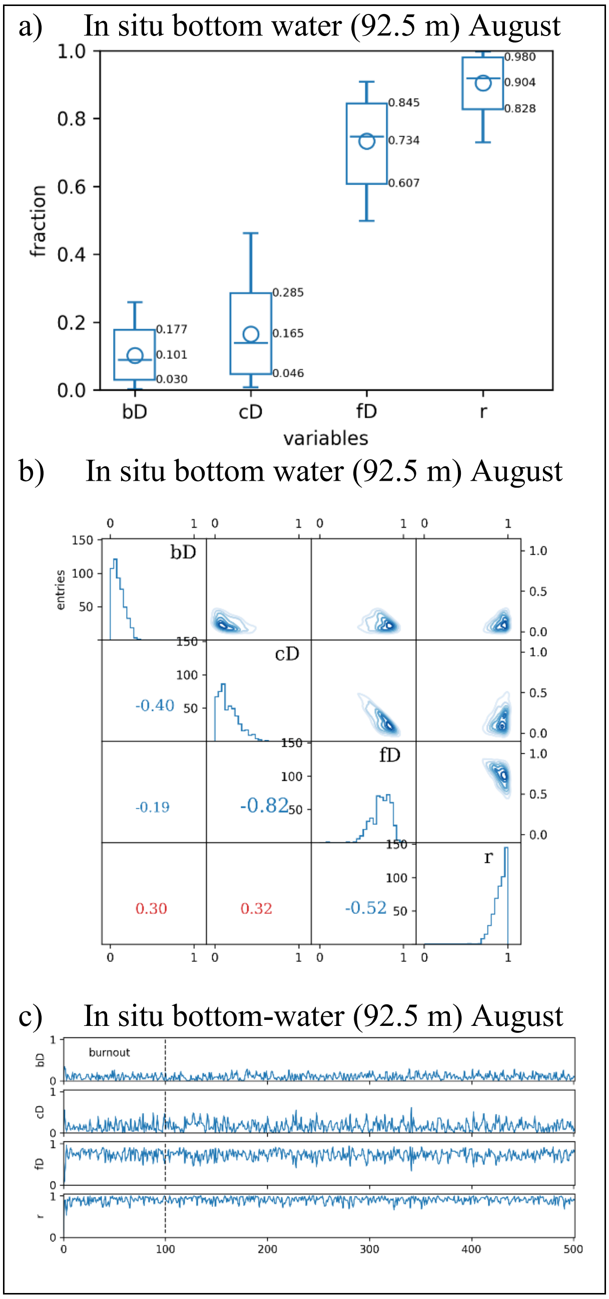


**Fig. S5**: 2D FRAME model output for the in situ bottom water sample (92.5 m, August). Panels show: (**a**) Summary plot of distributions with key statistics (mean = circle, median = horizontal line, 68% confidence interval (CI) = box, 95% CI = whiskers); (**b**) Histograms showing the probability distributions of mixing fractions (diagonal), with correlation displayed as contour plots (upper right), and numerical correlation coefficients (lower left); (**c**) Markov chain illustrating model iteration progress, including mixing configurations and the number of iterations needed to reach stabilization. Only data after the point of stabilization (dashed vertical line) are used, and iterations before are discarded, and marked as burnout.

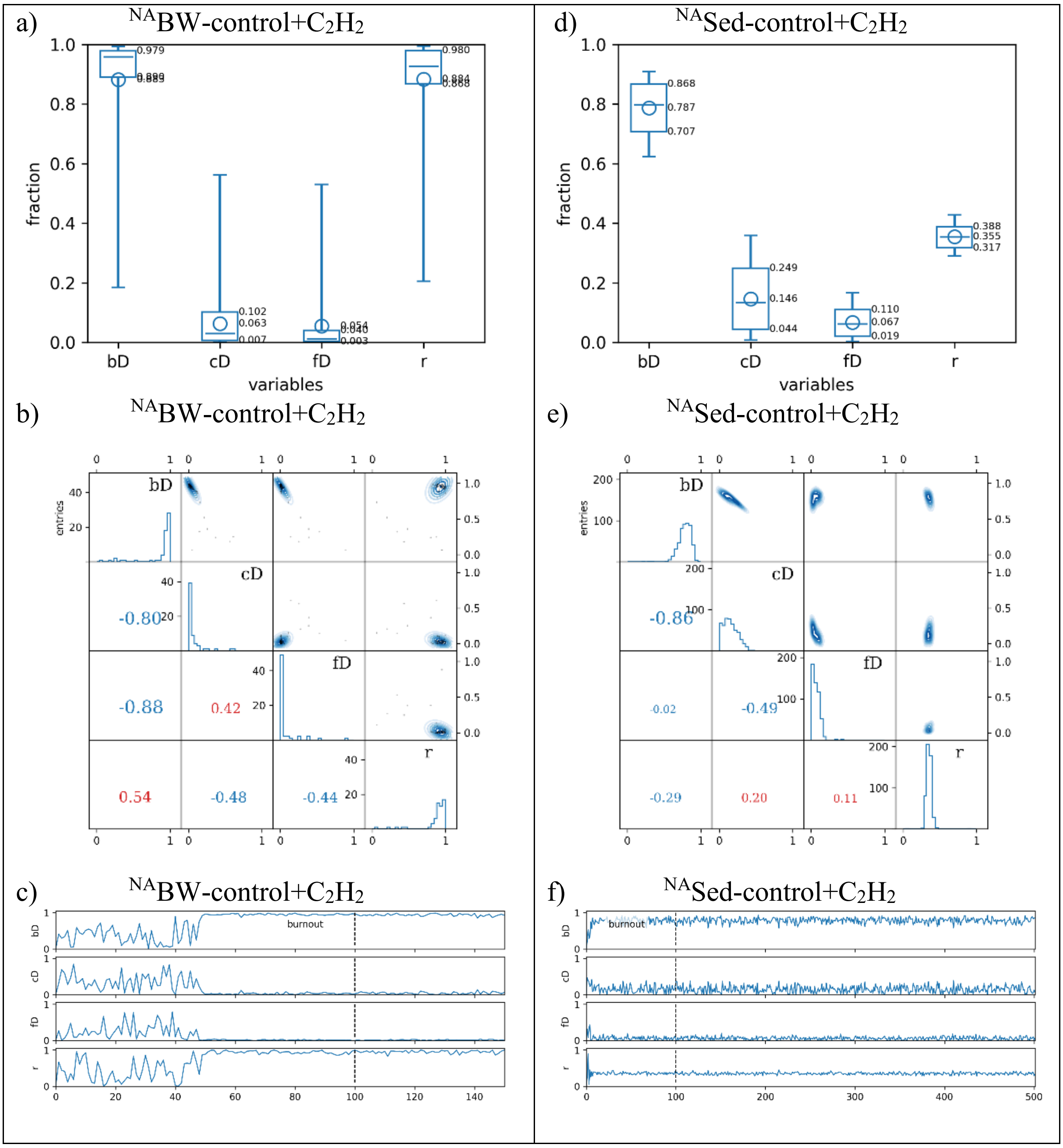


**Fig. S6**: 3D FRAME model output for $^{NA}$BW-control+$C_2H_2$ and $^{NA}$Sed-control+$C_2H_2$ samples. Panels show: (**a, b**) Summary plot of distributions with key statistics (mean = circle, median = horizontal line, 68% confidence interval (CI) = box, 95% CI = whiskers); (**c, d**) Histograms showing the probability distributions of mixing fractions (diagonal), with correlation displayed as contour plots (upper right), and numerical correlation coefficients (lower left); (**e, f**) Markov chain illustrating model iteration progress, including mixing configurations and the number of iterations needed to reach stabilization. Only data after the point of stabilization (dashed vertical line) are used, and iterations before are discarded, and marked as burnout.

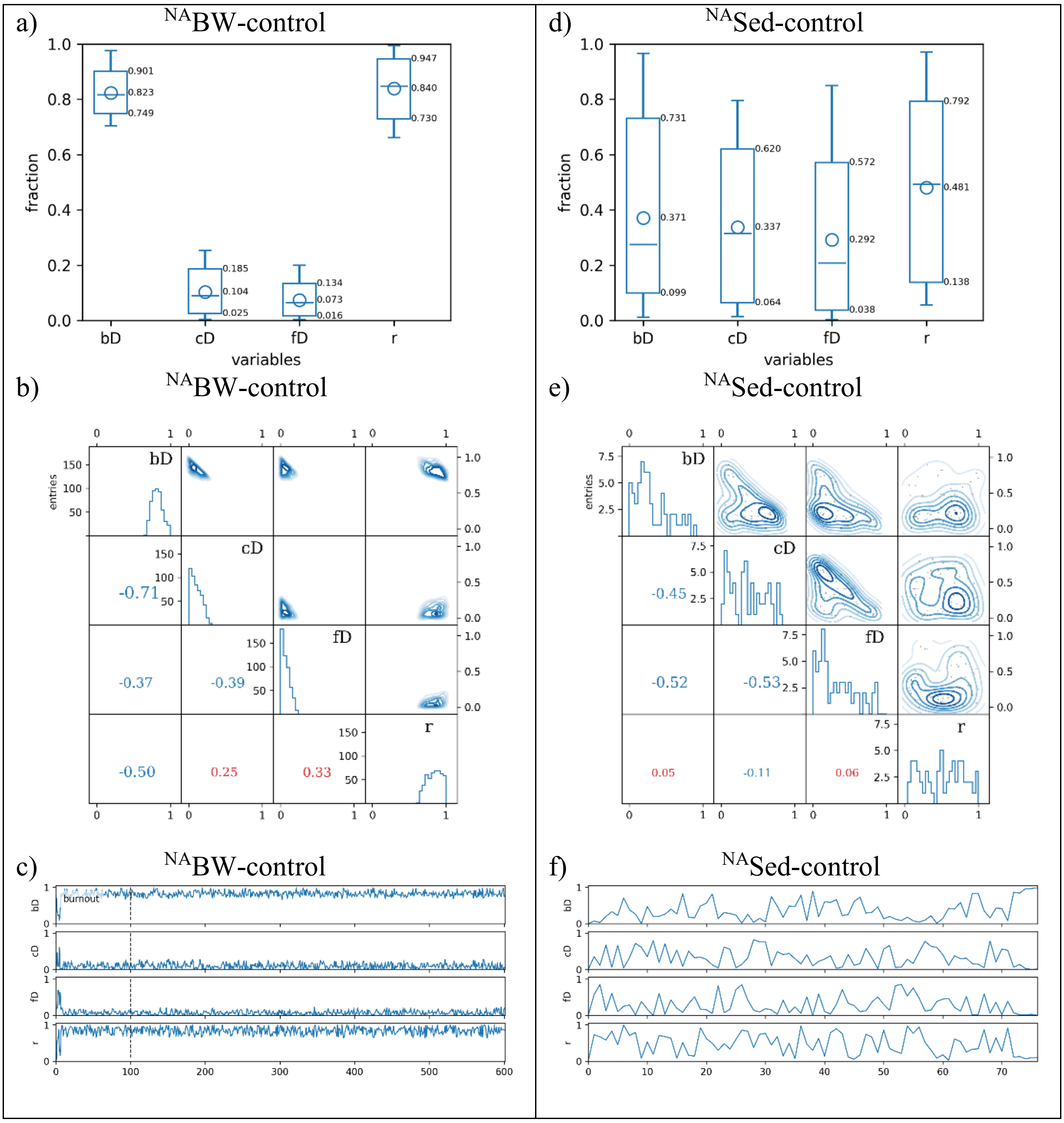


**Fig. S7**: 3D FRAME model output for $^{NA}$BW-control and $^{NA}$Sed-control samples. Panels show: (**a, b**) Summary plot of distributions with key statistics (mean = circle, median = horizontal line, 68% confidence interval (CI) = box, 95% CI = whiskers); (**c, d**) Histograms showing the probability distributions of mixing fractions (diagonal), with correlation displayed as contour plots (upper right), and numerical correlation coefficients (lower left); (**e, f**) Markov chain illustrating model iteration progress, including mixing configurations and the number of iterations needed to reach stabilization. Only data after the point of stabilization (dashed vertical line) are used, and iterations before are discarded, and marked as burnout. No burnout was reached for the $^{NA}$Sed-control.

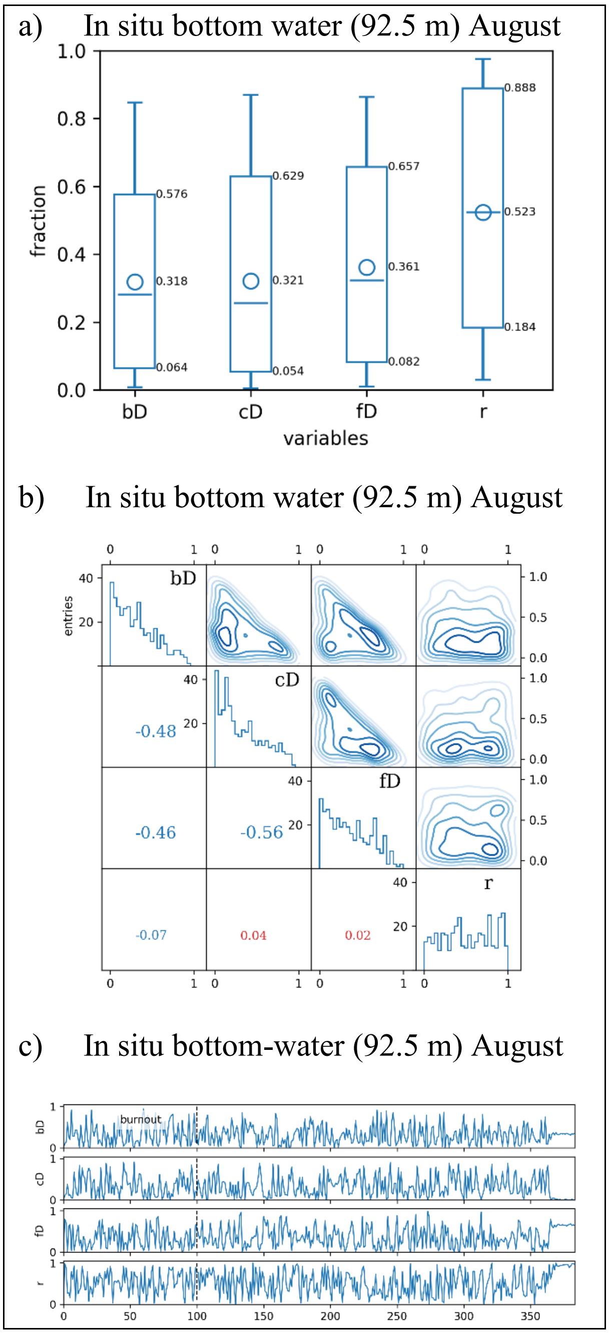


**Fig. S8**: 3D FRAME model output for the in situ bottom water sample (92.5 m, August). Panels show: (**a**) Summary plot of distributions with key statistics (mean = circle, median = horizontal line, 68% confidence interval (CI) = box, 95% CI = whiskers); (**b**) Histograms showing the probability distributions of mixing fractions (diagonal), with correlation displayed as contour plots (upper right), and numerical correlation coefficients (lower left); (**c**) Markov chain illustrating model iteration progress, including mixing configurations and the number of iterations needed to reach stabilization. Only data after the point of stabilization (dashed vertical line) are used and iterations before are discarded, and marked as burnout. Large oscillations in the Markov chain indicate that the model could not identify a stable or probable solution.
